# A human developing heart atlas reveals cardiovascular cell diversity for in vitro model benchmarking

**DOI:** 10.1101/2024.04.27.591127

**Authors:** Vincent Knight-Schrijver, Semih Bayraktar, James Cranley, Kazumasa Kanemaru, Beatrice Waller, Maria Colzani, Chunk-Ka Wong, Hongorzul Davaapil, Jonathan Chuo Min Lee, Maeve D’Souza, Maria Katharina Köhne, Krzysztof Polanski, Laura Richardson, Claudia I. Semprich, Rakeshlal Kapuge, Monika Dabrowska, Ilaria Mulas, Shani Perera, Minal Patel, Siew Yen Ho, Xiaoling He, Hung-Fat Tse, Richard Tyser, Laure Gambardella, Sarah A. Teichmann, Sanjay Sinha

**Author notes:** Co-first authors (Vincent Knight-Schrijver, Semih Bayraktar, James Cranley, Kazumasa Kanemaru).

## Abstract

The human heart and adjoining great vessels consist of multiple cell types essential for life, yet many remain uncharacterised molecularly during development. Here, we performed a high-resolution profiling of the developing heart and great vessels between 4 and 20 post-conception weeks using single-cell and spatial transcriptomics defining 63 cell types with distinct identity and location-specific signatures. We reveal previously unreported molecular identities in cell types, including the pericardium and the ductus arteriosus. In the cardiomyocytes, we identify signatures of the trabeculated-compact, and right-left axes of ventricular cardiomyocytes. In vessels, we distinguish the constituents belonging to either coronary or great vessels. We confirm our transcriptional findings spatially, revealing nuanced signatures with specific zonation patterns and validating this atlas as a curated transcriptional reference for future studies. We leverage the temporal scope of the presented atlas to build `CMageR`, a predictive pipeline for scRNA-seq combining cardiac cell annotation with a transcriptional cardiac clock of single-cell developmental age for each cell type. Our cardiomyocyte clock captures dynamic biology, revealing core functional changes and novel markers of maturity during the first and second trimester. Finally, we benchmark in vitro models, suggesting a transcriptional right-chamber bias in stem cell derived cardiomyocytes with the oldest model age-matched to 12 post-conception weeks. Collectively, our work provides a high-resolution atlas of human cardiac development to enhance our understanding of function in development, health, and disease, and a foundation for building a rich reference to benchmark and improve in vitro models.

## 1 Introduction

The heart is the first organ to function during embryonic development, where successfull cardiogenesis requires the differentiation and coordination of multiple different cell types. The heart and vessels are also major clinical targets, as cardiac diseases are the primary component of global disease burden in adults. As such, the heart is a primary focus of stem cells, with platforms that focus on modelling cardiogenesis, regenerative medicine, and new approach methodologies (NAMs). However, many of these models are currently developed in isolation from the reality of the native developmental environment, often resulting in cell states and developmental ages that are disconnected from their intended translational use. A deeper understanding of the transcriptomic and epigenomic landscape of the unique cellular constituents of the developing human heart is critical, helping us understand how these cells coordinate during development and how this can go awry to cause disease, congenital or adult-onset cardiac complications, or how faithfully *in vitro* models capture the developmental cardiovascular cell diversity and ageing processes. To fully understand the cellular and molecular mechanisms at play during this cardiogenesis, we must first define the repertoire of cell types in the developing heart.

With the advent of single-cell transcriptomic and epigenomic technologies, alongside spatial transcriptomics, many researchers have leveraged the mapping of cellular signatures to better understand organogenesis, including the developing human heart[1–8]. While these studies offered valuable insights into cardiac development, several cell types are still unresolved at high-resolution due to factors including a limited number of samples, inadequate coverage of developmental stages (focusing on either the 1st or 2nd trimester), or limited depth of spatial localization.

In this paper, and in our accompanying paper[9], we aimed to address these limitations by providing a high-resolution spatio-temporal and multi-omic atlas of the developing human heart and great vessels. Consequently, we performed single-cell RNA-sequencing using 21 hearts across the first and second trimesters[9]. In this paper, we establish the diverse array of cell types that make up the first and second trimester human heart and shed light into previously unreported cell types, including the cells of the pericardium or the ductus arteriosus. We identify distinct signatures that constitute the left-right, and compact-trabeculated axes of cardiomyocytes (CMs). Within the conduction system, we resolved the sinoatrial node and atrioventricular node pacemaker cells, alongside proximal and distal cardiac conduction system cells. Through targeted dissection and sequencing of samples, we differentiated the signatures between similar cellular constituents across coronary and great vessels and identified enhanced Notch signalling within coronary smooth muscle cells compared to their great vessel counterparts. Our analysis indicates that cells of the great vessels collectively are enhanced in synaptic assembly. We infer increased prostaglandin signalling, alongside reduced endothelin sensing, at ductus arteriosus smooth muscle cells to maintain the patency of ductus arteriosus during development.

Lastly, alongside additional datasets, we packaged our atlas into a trained and validated proof-of-concept computational reference pipeline for predicting healthy developmental cardiovascular cell diversity and developmental age in single-cell RNA sequencing data (`CMageR`). Using this, we deeply characterise published stem cell-derived cardiomyocyte (hPSC-CM) models, revealing a range of different CM purities, CM subtypes, and developmental ages in current protocols, as well as a right-chamber transcriptional bias found broadly across *in vitro* models of ventricular CMs.

## 2 Results

To investigate cellular composition, heterogeneity, and interactions in the developing human heart, with our accompanying paper[9], we collected a total of 21 human fetal hearts between the ages of 4 and 20 post-conception weeks (PCW) (12 female, 9 male, (**Supp. Fig. S1a**) for single-cell RNA or single-nucleus RNA sequencing (Fig. 1a). The hearts were small enough to process whole, enabling us to capture the full breadth of cellular heterogeneity, including rare populations, and explore cell types at different stages of maturity. To gain regional insight, some of these hearts were dissected into several distinct anatomical regions, such as node, great vessels, ductus arteriosus, or pericardium, prior to sequencing (Fig. 1a).

**Fig. 1:**
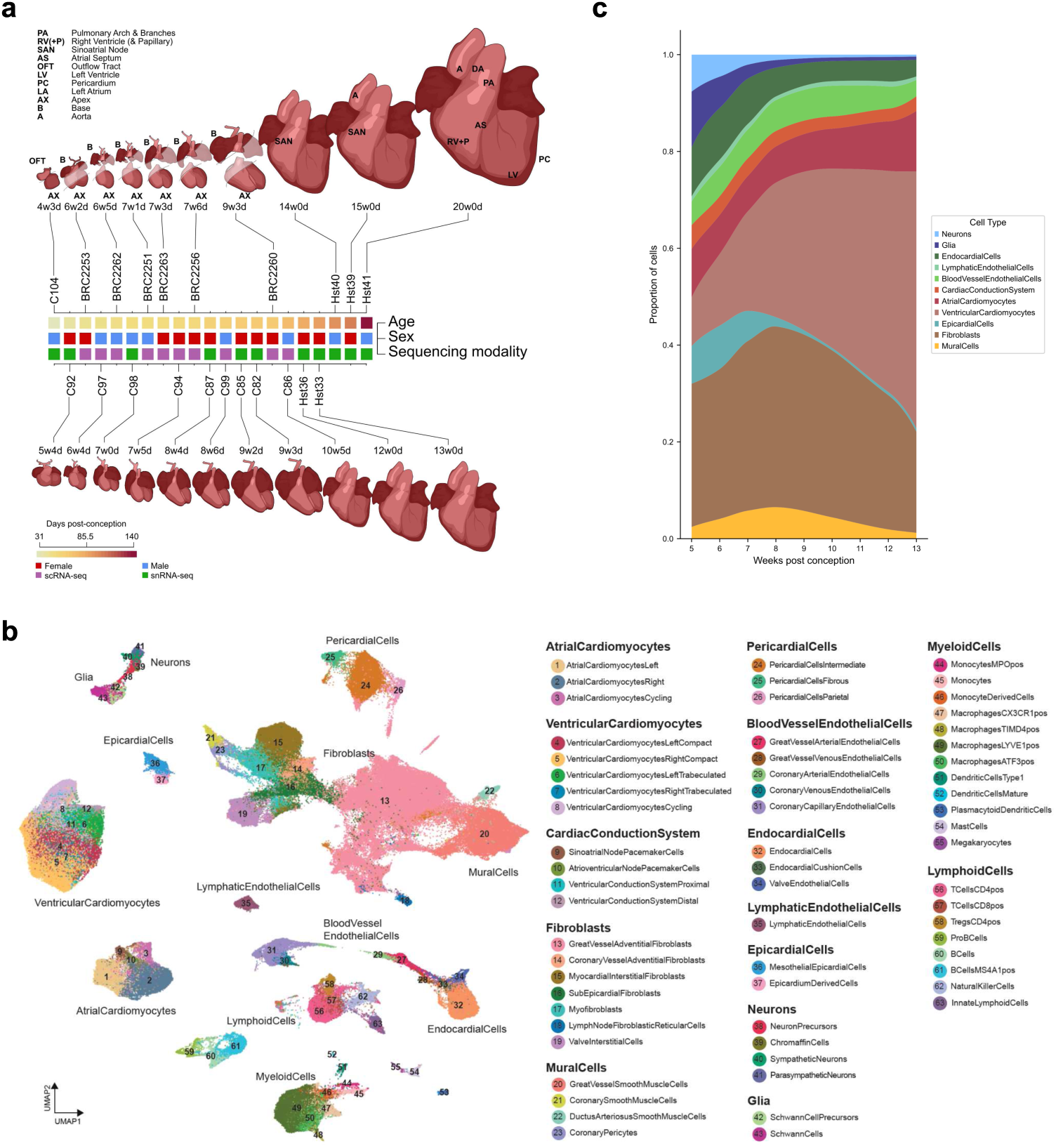
Atlas of the developing human heart and the great vessels. **A:** Overview of study design and data modalities. Single-cell or single-nuclear data were generated from a total of 21 human fetal hearts between the ages of 4 and 20 PCW. Targeted dissection was performed on the upper row samples as indicated. Lower row hearts were sampled in entirety. **B:** Two-dimensional uniform manifold approximation and projection (UMAP) embeddings of gene expression of 63 cell types of the developing human heart and the great vessels, spread across 14 mid- and 6 coarse-grains. Fine-grained cell type annotations are provided adjacent to the UMAP embeddings, grouped under their respective mid-grains. **C:** Changes in the composition of the heart in the first trimester, depicted as proportion. To minimize sampling bias, immune cells (enriched by CD45 sorting), great vessel cells and pericardium were excluded, as these were inconsistently represented across samples. Only hearts processed in entirety, rather than by targeted dissection, were included in this analysis.

After quality control, 297 473 cells and nuclei were retained for further analysis. Cell type annotation was performed in gene expression space, with batch correction using scVI (**Supp. Fig. S1**)[10, 11]. A coarse-grained annotation identified 6 major cell types representing Cardiomyocytes, Endothelium, Epicardium, Mesenchymal cells, Leukocytes and Neural cells. Iterative rounds of annotation revealed 14 mid-grained and 63 fine-grained cell types. Wherever possible, cells were annotated to match a specific function or anatomical location, such as the coronary and great vessel constituents (**Fig. 1b, S2a**). No adipocytes were detected in our samples, in agreement with previous work showing they are first observed in the third trimester of development and later than we sampled[12]. The comprehensive lists of fine grain cell numbers by donor, region and post-conception week are tabulated in Supplementary Table 2.

Using these annotations, we assessed the changing cellular composition across time within only the whole heart nuclei RNA-seq samples. Our main observation was a decrease in the relative abundance of non-myocyte populations coupled with a rise in the fraction of CMs, especially those of the compact ventricular myocardium (**Fig. 1c, S2d, e**). This suggests that the later stages of the first trimester are important for the expansion of compact myocardium.

### 2.1 Cardiomyocyte diversity in development

Subclustering of Cardiomyocytes (CMs) yielded 12 fine-grain CM subtypes, grouped into three mid-grained cell types based on anatomy function; AtrialCardiomyocytes and VentricularCardiomyocytes (aCM and vCM), and CardiacConductionSystem (CCS) cells (**Fig. 2a**). Using data from Cranley et al.[9] these populations were validated through spatial transcriptomics by mapping their transcriptional profiles to their corresponding positions using cell2location. This highlighted the distinct localisation of atrial and vCM subpopulations (left (L), right (R), compact and trabeculated) (**Fig. 2b**), and confirmed the position of the CCS subpopulations (**Fig. 2c**). Overall, the well-established cardiac gene *TNNT2* was the most sensitive CM marker in the developing heart, regardless of the subtype of CMs (**Supp. Fig. S4**), and a gradual decrease was seen in the proportion of cycling cells from earliest to latest timepoints (**Fig. 2d**).

**Fig. 2:**
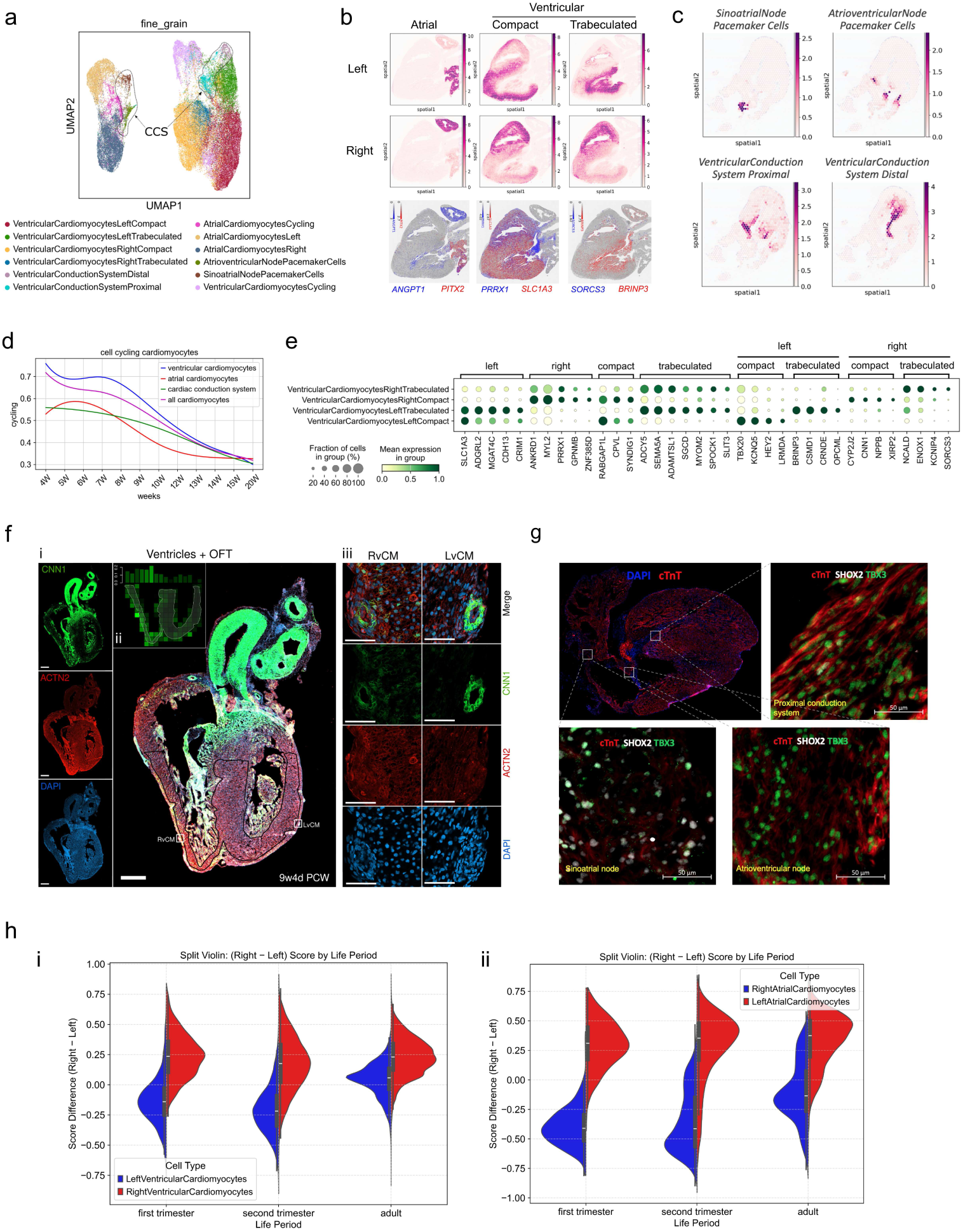
Cardiomyocyte diversity in the developing heart. **A:** Two-dimensional uniform manifold approximation and projection (UMAP) embeddings of cardiomyocytes (CMs) using the coarse grain subset of CMs-only displaying fine grain cell type labels. CMs from all stages were used to generate this UMAP. CCS=cardiac conduction system. **B:** Spatial mapping of working CMs onto a visium spatial transcriptomics section from a human 16 post-conception week (PCW) heart using cell2location with selected left or right markers for atrial CM, compact ventricular CM, or trabeculated ventricular CM markers. **C:** Spatial mapping of CCS CMs onto a visium spatial transcriptomics section from a human 7 PCW heart using cell2location. **D:** A decreasing proportion of cycling CMs seen across mid grain annotations, indicating a reduction in cycling CM abundance through development. **E:** Differential expression analysis results between ventricular CMs as shown in a dot plot depicts the top transcriptional markers for left or right, trabeculated or compact ventricular CM subtypes. **F:** Immunofluorescence validation of CNN1 in a human fetal heart at 9 weeks and 4 days postconception showing higher CNN1 expression in right ventricular compact cardiomyocytes in i, the whole section; ii, quantified as a significant monotonic decrease in expression moving from right to left (Spearman’s ranked correlation, rho = 0.75, p *<* 0.001); and iii, compared with CNN1 positive smooth muscle cells of a coronary artery at higher magnification. Scale bars are 500 µm (10x magnification, whole heart), and 50 µm (40x magnification, selected regions). **G:** Immunofluorescence validation of the CCS. TBX3 expression is observed at Sinoatrial Node (SAN) pacemaker ells, Atrioventricular Node (AVN) pacemaker cells and proximal CCS, while SHOX2 expression is observed at SAN pacemaker cells only. **H:** Violin plots showing right-left score differences across life stages in atrial and ventricular cardiomyocytes. At each period, a right and left score was calculated between the cell types using the right / left differentially expressed genes (DEGs) identified between the right and left chamber cells at that life period. The difference of right and left scores are displayed as split violin plots showing that aCM transcriptional chamber differences continue in adult hearts while vCM transcriptional chamber differences converge.

The top markers for CM subpopulations were identified using differential expression analysis. In the atria, *PITX2*, a regulator of left-right asymmetry associated with the left atrium[13], was a specific marker of left aCMs (**Fig. S2b**, **Supp. Fig. S5a**). *PANCR*, a *PITX2* -adjacent long non-coding RNA which has been shown to positively regulate *PITX2* expression[14], was highly specific to left aCMs as well, underlining the critical role of *PITX2* in LaCM cell identity. Alongside the previously reported *BMP10*, the RaCMs expressed *NTM* and *ANGPT1* (**Fig. S2b**, **Supp. Fig. S5a**). Right atrial-derived angiopoietin has been shown to be critical for coronary vein formation in mice[15], suggesting a similar role in humans.

In the ventricles, *SLC1A3* and *PRRX1* were differentially enriched in LvCMs and RvCMs respectively, corroborating current knowledge (**Fig. 2b**, **Fig. 2e**, **Supp. Fig. S5b**)[16]. Our dataset revealed additional markers such as *CDH13* in LvCMs, or *ANKRD1*, *MYL2*, *GPNMB*, and *CNN1* in RvCMs (**Fig. 2b**, **Fig. 2e**). Using available spatial transcriptomics data[16], the LvCM and RvCM cluster annotations were spatially validated while immunohistochemistry orthogonally confirmed CNN1 as a marker of compact RvCMs at protein level, showing a significant decrease in expression in left ventricles (LV) compared with right ventricles (RV) (p < 0.05, n = 3, Student’s t-test) (**Fig. 2b**, **Supp. Fig. S5c**). To test whether *PRRX1* was a key transcription factor regulating right vCM identity, a partial knockdown of *PRRX1* was performed in hPSC-CMs, resulting in a significant downregulation of *ANKRD1*, *CNN1*, and *CYP2J2* (**Supp. Fig. S5d**). Lastly, trabeculated CMs were separable from compact CMs in both ventricles with the expression of *SGCD*, *SPOCK1* and *SLIT3*, highlighting roles in membrane stability and ECM remodelling in trabeculated vCMs, and with *BRINP3*, and *SORCS3* more specifically in left or right trabeculated vCMs respectively (**Fig. 2b**, **Fig. 2e**).

The CCS cluster did not have any marker that could capture all its constituents in a specific manner; *CNTN2*, *NPTN* and *NTM*, used to mark the entire mouse CCS[17], did not exhibit the same specificity in the developing human heart (**Supp. Fig. S5f**). However, subclusters within the CCS were identifiable. *MYH6*, *CACNA1D*, *NR2F2* and *TBX3* jointly marked the sinoatrial node (SAN) and atrioventricular node (AVN) pacemaker cells, whereas *MYH7*, *NAV1* and *LPL* marked the ventricular conduction system (VCS) cells. The marker profile and band-like localisation of the AVN mapping at ventricles and atria suggested that these cells could be part of a primitive atrioventricular (AV) myocardium. *SHOX2*, *PDE1A* and *TENM4* were specific for SAN pacemaker cells, whereas *GREB1L*, *NRXN3* and *RSPO* were specific to AVN pacemaker cells (**Supp. Fig. S5f**). We validated *SHOX2* specificity at a protein level using immunohistochemistry showing SHOX2+ and TBX3+ SAN compared with a TBX3+ but SHOX2AVN (**Fig. 2g**). Additionally, *MYH11*, one of the most specific markers of smooth muscle cells, was expressed at the SAN pacemaker cells, in line with previous reports [18, 19], and *NIBAN1* and *PHACTR1* were expressed by proximal VCS while *OPCML*, *NTN1* and *IRX2* marked distal VCS cells (**Supp. Fig. S5f**).

To determine if the LvCM or RvCMs chamber-specific transcriptional signatures were time-associated, we compared their overall signatures throughout fetal stages, and against their adult counterparts using published data[18]. Differential expression analyses between LvCMs and RvCMs and scoring of overall L / R signatures at either first trimester, second trimester, or broad adult stage provides some evidence that chamber transcriptional identities partially converge throughout the human lifespan (**Fig. 2h**, **Supp. Fig. S6**). This was further noted as many chamber-specific genes, including *PRRX1*, *SLC1A3*, and *CNN1* were differentially trending but not significant between RvCM and LvCM cells in adult hearts (**Supp. Fig. S6**). Applying the same approach to aCMs revealed a similar trend. However, the chamber specificity scores in aCMs were less similar than those of the vCMs (**Fig. 2h**), and many LaCM or RaCM specific genes were still different in adult hearts (**Supp. Fig. S6**).

Our combination of single cell, spatial transcriptomics, and orthogonal protein-level validation of chamber-specific transcriptional signatures and key markers provides strong evidence that we have captured and annotated the CMs from all four chambers of the developing human heart and its conduction system. If these differences persist in adults ventricles, they may do so only mildly.

### 2.2 Cell diversity of coronary and great vessels

Both mural cells and endothelial cells (ECs) within the coronary and great vessels are known to have multiple embryological origins[20, 21]. These distinct origins along with specific local cues give rise to a wide range of differing vascular cell states that are poorly characterised in human development yet have important roles in vascular development and function.

To investigate this cardiovascular diversity in more detail, we examined the endothelium, fibroblasts and smooth muscle cells (SMCs) collected at either apex or base dissociated samples, enabling us to identify these cell types based on regional distribution. Through our targeted dissection strategy (**Fig. 1a**), we could identify coronary SMCs (CSMCs) from their localisation to both apex and base, whereas great vessel SMCs were observed only at the base (**Fig. 3a**). Using the same framework, we distinguished the endothelial and fibroblast constituents of the coronary and great vessels (**Fig. 3b**), which was supported by spatial transcriptomics[9], where we observed a clear separation in the mapping of relevant cell types across spatial slides (**Fig. 3c**). This molecular differences between great and coronary vessels to be revealed. For example, great vessel adventitial fibroblasts expressed *ROR1* and the mechanosensing ion channel *PIEZO2*, while coronary vessel adventitial fibroblasts expressed *TBX20* and *TCF21* (**Fig. 3d**). We were interested in identifying vessel-wide biological functions shared across the constituents of either vessel type. Overall, ECs, mural cells and fibroblasts of the great vessels shared significant Gene Ontology biological process (GO:BP) terms relating to neurogenesis, sensory organ development or axon guidance. All these cellular components expressed *NLGN1*, with a potential role in synaptic assembly and guiding the formation of the cardiac plexus, the network of autonomic nerve fibres and ganglia, that sits on the great vessels (**Fig. 3e**, **Supp. Fig. S7a**).

**Fig. 3:**
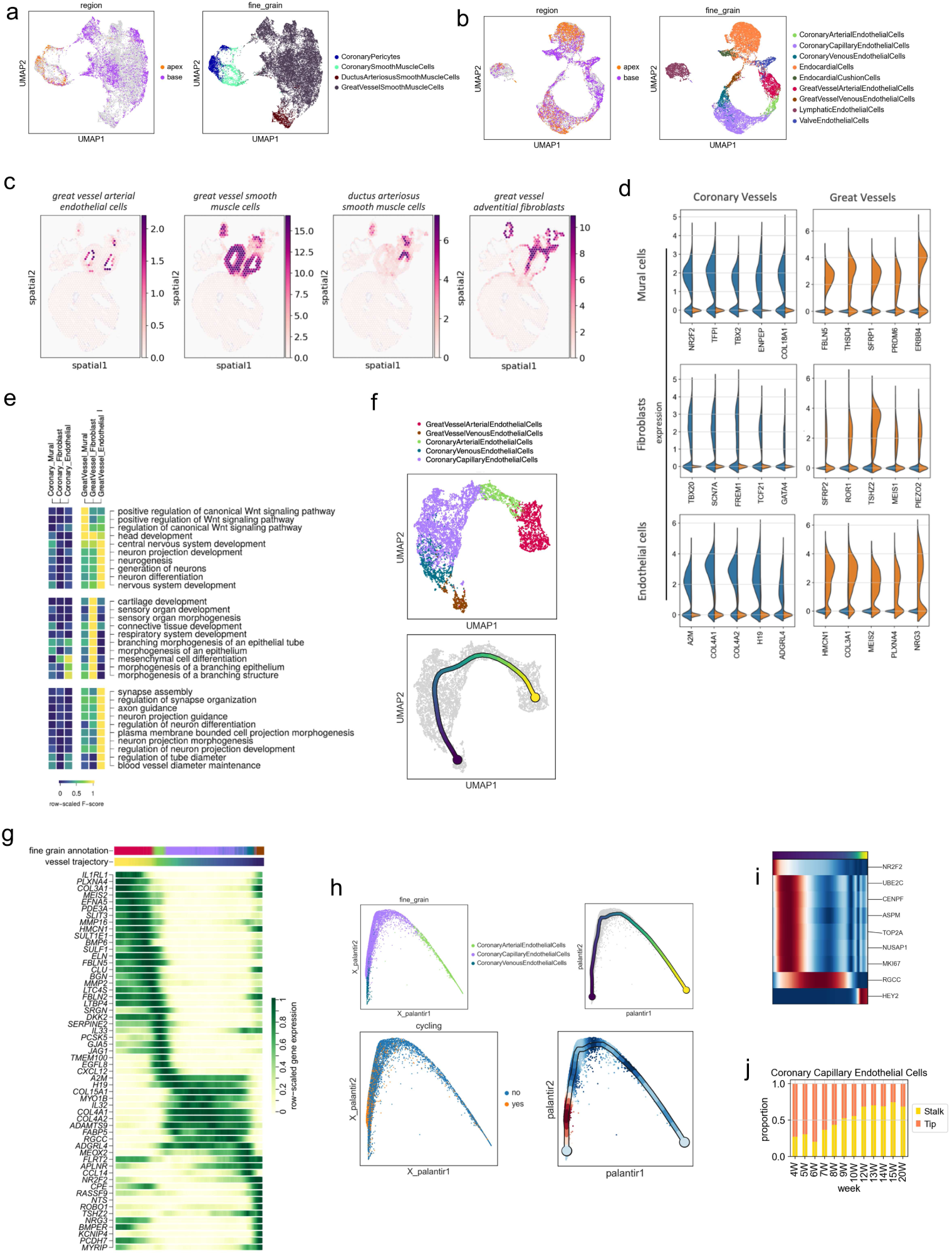
Profiling of coronary and great vessel identities within the developing human heart. **A, B:** Two-dimensional uniform manifold approximation and projection (UMAP) embeddings made from a, smooth muscle cell (SMC) cell types and b, endothelial cell types from all samples and stages. Cells sourced from apex and base dissection regions in samples were highlighted in the left panel in each case. For example, coronary SMCs are expected in both apex and base sourced samples whereas great vessel SMCs are expected in only base sourced samples. Cells from other regions in these region panels are coloured in grey. **C:** Spatial mapping of great vessel constituents; mapping absent from myocardium, highlighting the high-resolution nature of annotations and the dataset. **D:** Split violin plots for the expression of coronary (blue) vs great vessel (orange) enriched genes in vessel constituents. y-axis indicates log-expression of normalised counts. **E:** Gene Ontology analysis results showing top-scoring significant terms for all coronary and great vessel constituents, suggesting that great vessel constituents have shared terms relating to neural processes. F-scores were calculated using the query size, intersection size, and domain size and then transformed within each term linearly by subtracting the minimal F from all celltype subsets, and dividing by the new maximum F. This standardisation approach highlights within-term, not between-term, enrichment across the cell types. **F, G:** Two-dimensional UMAP embeddings made from cells sourced from all stages depicting f the construction of endothelial continuum of coronary circulation based on vascular anatomy and flow with g, the expression profile of fine-grain blood vessel endothelial cells along this continuum. **H, I:** Embeddings of coronary ECs, presented with fine grain label, trajectory, and cell cycling information as h, the continuum of coronary vessel endothelial cells with individual UMAPs depicting fine grain, pseudotime, cycling cell and cell cycle score embeddings along the continuum; and i, the expression of cell cycling modules along the continuum with venous but not arterial cells in proximity to cycling modules.

We identified great vessel and coronary ECs with arterial, venous and capillary specifications, which appeared to align sequentially with the vessel size, function, and blood flow (**Fig. 3b**). To better capture this, we used trajectory analysis to form a continuum of the ECs present within the coronary circulation as the vessels make their way through the heart, highlighting the molecular profiles that vessel-lining ECs take on as a result of the changing environments (**Fig. 3f**). For example, *HMCN1* and *PLXNA4* were expressed by the ECs of the great vessels, while coronary vessel ECs expressed *A2M* and *ADGRL4* (**Fig. 3g**). It has been suggested that signalling through *NR2F2*, expressed in the venous clusters here, blocks pre-arterial specification and activates cell cycling genes[22, 23]. We observed that cycling cells of the developing coronary endothelium have a closer transcriptional identity to the venous and capillary coronary ECs (**Fig. 3h, i**). Sprouting angiogenesis is achieved by endothelial tip cells, and within the capillary coronary endothelium, we observed the tip identity peaked around 6 PCW and then gradually decreased through our time points (**Fig. 3j**).

### 2.3 Mural cell diversity

We observed transcriptional differences in the SMC types and validated our finding with immunohistochemistry. For example, coronary and great vessel SMCs expressed PLA2G5 and FOXC1, respectively, in a mutually exclusive manner (**Fig. 4a**). Compared within coronary mural cells, coronary pericytes expressed *THBS4*, *KCNJ8* and *CYGB*, whereas CSMCs expressed *ELN*, *TPM1* and *ACTA2* (**Supp. Fig. S7b**). Our most mature sample (20 PCW) allowed us to confidently dissect the aorta and pulmonary artery during sampling and sequence separately. We observed a pulmonary specification within pulmonary artery SMCs with enriched expression of *INPP4B*, *PXDNL* and *ALDH1A2* (**Supp. Fig. S7c**).

**Fig. 4:**
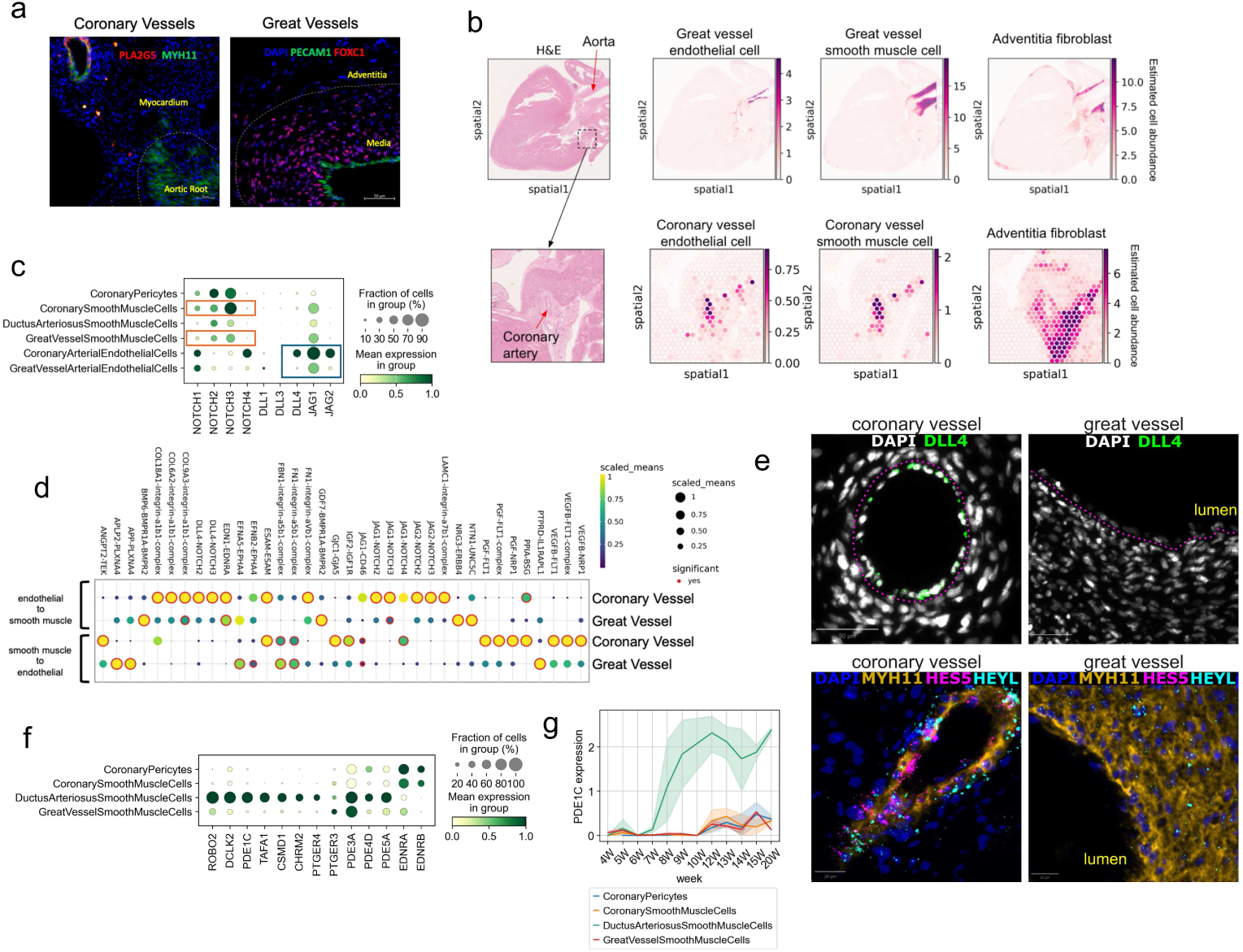
Mural Cell Diversity. **A:** Validation of distinct expression of PLA2G5 and FOXC1 (protein, immunohistochemistry) at Coronary smooth muscle cells (CSMC) and great vessel SMCs (GVSMCs). Scale bars, 50 µM. **B:** Mapping of vessel constituents across spatial transcriptomics sections underlines the indepth annotation of the atlas. First row, great vessel constituents. Second row, coronary vessel constituents. **C:** Dot plot depicting the expression of Notch receptors and ligands among mural and arterial endothelial cells in the dataset. CSMCs and GVSMCs exhibit a similar Notch receptor profile (orange boxes), while their respective arterial endothelial cells (blue box) exhibit differential expression of ligands, with *DLL4* and *JAG2* being specifically expressed at coronary arterial ECs. **D:** The interactions between ECs and SMCs of the coronary and great vessels at the intima-media interface, despite sharing similar cell types, exhibit distinct interactions. **E:** RNAscope and protein immunohistochemistry validation of differential Notch signalling at coronary vessels. The top panel shows RNAscope detection of *DLL4* specifically in coronary arterial ECs. The bottom panel shows increased *HEYL* expression and specific *HES5* expression in CSMCs detected by RNAscope, visualised on the same slides using *MYH11* immunohistochemistry to mark SMCs. **F:** Differentially expressed markers of DASMCs in comparison to the other mural cells in the atlas. **G:** Temporal expression pattern of *PDE1C* across mural cells.

Across mural cells, we observed differential expression of canonical Notch downstream effectors, including *HES5*, *HEY2*, and *HEYL* in CSMCs (**Supp. Fig. S7d**). While SMCs from both vessels displayed a comparable profile of Notch receptor expression, the ligands provided by their arterial EC counterparts differed markedly (**Fig. 4c**). Investigating cell to cell communication mechanisms at both the vessel types, we observed a pronounced Notch signalling towards coronary vessel SMCs, mediated by *JAG1*, *JAG2* and *DLL4* at intima-media interface (**Fig. 4d**), and by *DLK1* at media-adventitia interface (**Supp. Fig. S7e**). In particular coronary arterial ECs expressed *DLL4* and *JAG2*, with *DLL4* -*NOTCH3* interaction, a ligand-receptor pair known for potent activation of *NOTCH3*, emerging as a strong candidate for driving the heightened activation of Notch downstream signalling and downstream transcription factor expression (**Fig. 4e**).

Closure of the ductus arteriosus is essential immediately after birth to ensure proper cardio-pulmonary functioning, as a patent ductus arteriosus (PDA) may lead to respiratory distress and heart failure. We identified the SMCs of the ductus arteriosus (DASMCs) with a distinct expression profile of *DCLK2*, *PDE1C* and *TAFA1* compared to other mural cells in the dataset (**Fig. 4f**). The expression of PDE1C increases sharply around 7 weeks into development (**Fig. 4g**) and given it encodes for a phosphodiesterase, this might be implicated in the vasoconstrictor function of the ductus arteriosus. We observed a specific expression of prostaglandin receptor *PTGER4* and muscarinic M2 receptor *CHRM2* in DASMCs, with a reduced endothelin receptor expression (*EDNRA* and *EDNRB* ; (**Fig. 4f**)) when compared to other mural cells in the dataset. PTGER4 has been shown to play a role in ductus arteriosus closure in mice[24]. Investigating the ECs obtained from the ductus arteriosus sampling, these cells had an enriched expression of *PDE4D*, alongside genes implicated in prostaglandin synthesis (**Supp. Fig. S7f**). Collectively, these suggest a mechanism whereby the duct is primed for closure, yet is kept open due to active prostaglandin signalling and reduced endothelin sensing during development.

### 2.4 Pericardium and epicardium

The pericardial and epicardial layers remain relatively poorly studied in terms of cellular composition and function despite their essential roles in cardiac development. We performed a targeted dissection of the pericardium from one donated heart (Hst41 20 PCW, (**Fig. 1a**)) enabling definition of cell clusters corresponding to both fibrous and serous pericardial layers, including the parietal and visceral layers of the folded serous pericardium (**Supp. Fig. S8a**).

Firstly, we found the two previously described mesothelial epicardium or migratory epicardium-derived cell (EPDC) populations of epicardial cells that make up the inner visceral layer of the serous pericardium (**Supp. Fig. S8a, b**)[7]. Compared with the other layers of the pericardium, both epicardial populations selectively expressed *ALDH1A2* and *EZR*. These cells also expressed the lubricant protein-coding gene *PRG4*, and hyaluronic acid synthesis gene *HAS1* (**Supp. Fig. S8b**), and interestingly, the interaction between lubricin and hyaluronic acid has been reported to synergistically enhance anti-adhesive properties[25]. Thus, epicardium-provided lubrication is implicated in reducing friction and minimising adhesions. Out of the common epicardial markers studied in animals and stem cell models, *BNC1* was seen to specifically define both populations of epicardial cells and was the most sensitive and specific marker of epicardial cells across the atlas (**Supp. Fig. S4**). *WT1* was expressed by the cells of the serous pericardium, including epicardium and controversially, *TCF21* expression was not relatively high in any of the epicardial or pericardial cells and was preferentially expressed in fibroblast clusters (**Supp. Fig. S4**). In addition to *TBX18*, the mesothelial epicardium was selective for *SBSPON*, and *ITLN1*, while the EPDCs were selective for *CPB1* and *VEGFC* (**Supp. Fig. S8b**). With evidence for VEGFC in driving angiogenesis[26], these cells potentially guide coronary vessel formation as they invade the developing myocardium to form coronary pericytes and SMCs.

Secondly, we found three clusters of cells describing either the parietal layer of the visceral pericardium, the fibrous pericardium, or a mixed cluster of pericardial cells with both visceral and fibrous markers (**Supp. Fig. S8a**). All three of these clusters expressed *COL6A3*, *COL12A1* and *COL16A1* (**Supp. Fig. S8b**). We found that the parietal layer of the serous pericardium distinctly expressed *TSHZ2*, *AUTS2* and *SETBP1*, while the fibrous pericardium expressed the chondrocytic marker *ITM2A* and procollagen peptidase enhancer *PCOLCE* (**Supp. Fig. S8b**). A strong expression of exosomal pseudo markers *CD9*, *CD63* and *CD81* was seen in fibrous and visceral serous pericardial cells (epicardium) (**Supp. Fig. S8b**), suggesting that these pericardial layers may contribute to the exosome rich pericardial fluid[27]. We spatially validated our annotations with immunohistochemistry using antibodies for TSHZ2 on a section of intact pericardium revealing TSHZ2 expression in non-myocardial tissue adjacent to the pericardial cavity, confirming these cells as parietal serous pericardium (**Supp. Fig. S8c**). Furthermore, cell2location mapping of the visceral serous pericardium onto spatial transcriptomics[9] showed an intramyocardial enrichment of EPDCs (**Supp. Fig. S8d**). Overall, these results reveal new markers of the pericardium suggesting functional differences between the pericardial cells (**Supp. Fig. S8e**). In particular, the expression of exosomal markers in some of these pericardial layers suggest they may represent the cellular source of the previously described exosome-rich pericardial fluid[27], implicating the pericardium as an active paracrine signalling compartment during cardiac development.

### 2.5 Endocardium and valves

The endocardium lines the interior of the heart chambers and, in addition to coronary endothelium, gives rise to the cardiac valves through a process of endothelial-to-mesenchymal transition (endo-MT) and endocardial cushion formation. Defects in this process are among the most common causes of congenital heart disease, meaning the molecular signals governing endo-MT and cushion cell identity have direct relevance to understanding both normal valve development and the origins of congenital septal and valvular defects. Within the endocardium and the valve lineage, cells of the endocardial cushion, valve endothelial and valve interstitial cells were identified, alongside endocardial cells (**Supp. Fig. S8f**). All the cells within this lineage expressed *POSTN* (**Supp. Fig. S8g**). *NPR3* expression was specific to the endocardium, where other endocardial markers *PCDH7* and *SMOC1* were shared with the endocardial cushion (**Supp. Fig. S8g**). Endocardial cushion cells on the other hand had a strong expression of *BMP4* and *HAS2*, and also expressed *PROX1*, *COL26A1*, *TMEM132D*, and *TSPAN8* (**Supp. Fig. S8g**). In fact, *TMEM132D* and *TSPAN8* emerged as specific markers of endocardial cushion cells across the atlas (**Supp. Fig. S4**). Endothelial cells of the valves express *APCDD1* along with endocardial cushion cells, however, had specific expression of *DKK2*, *KCNJ2* and *SULF1* within the endothelial lineage (**Supplementary Fig. 8g**). Valve interstitial cells on the other hand expressed *PTN*, *ID4* and *PRRX1*. These cells also expressed TGF-β signalling inhibitor *BAMBI*, together with endocardial cushion cells (**Supp. Fig. S8g**). Overall, *COL9A2* emerged as the most specific marker of valve interstitial cells across the whole dataset (**Supp. Fig. S4**). We were interested in cell-cell communication between myocardial layers and the epicardial or endocardial membranes the layers are nestled between. *ADGRG6* signalling, associated with the integrity of the compact wall and the identity of trabeculated CMs[28], originated from both myocardial layers towards the epicardium and the endocardium (**Supp. Fig. S8i, j, k**). Conversely, signals directed at *PLXNA4*, expressed by both compact and trabeculated CMs, originated from both the epicardium and the endocardium (**Supp. Fig. S8i, j**).

### 2.6 Leukocytes and Neurons

Within the coarse grain cluster of Leukocytes, we identified 20 sub-clusters at fine grain, encompassing a range of specific immune cell subsets across both myeloid and lymphoid lineages Additionally, coarse grain cluster of Neural cells was subclustered to 6 fine grain cell types including subsets of Neurons and glial cells. Our findings and categorisation of these cells and their markers are described in more detail in the supplementary results file (**Supp. Fig. S9**, **Supplementary Materials**).

### 2.7 A reference model of developmental cell diversity and age

We constructed and validated a single-cell transcriptional reference pipeline for cell type prediction with a cardiac biological clock for developmental age in the developing fetal heart (`CMageR`). The pipeline allows researchers predict the cardiovascular cell diversity and developmental age of query single-cell transcriptomics data, including from stem cell models. Our pipeline addresses prediction through two steps, firstly as cell type labelling through discrete classification using CellTypist[29], and secondly as a continuous model of cell type age using multivariate generalised additive models (mGAM) with the response variable as real chronological time (**Fig. 5a**). In both cases, efforts were made to carefully stratify the data, balancing the cell type composition among technical and biological variables such as sequencing modality and donor (**Supp. Fig. S10a**, **Supplementary Table 5**).

**Fig. 5:**
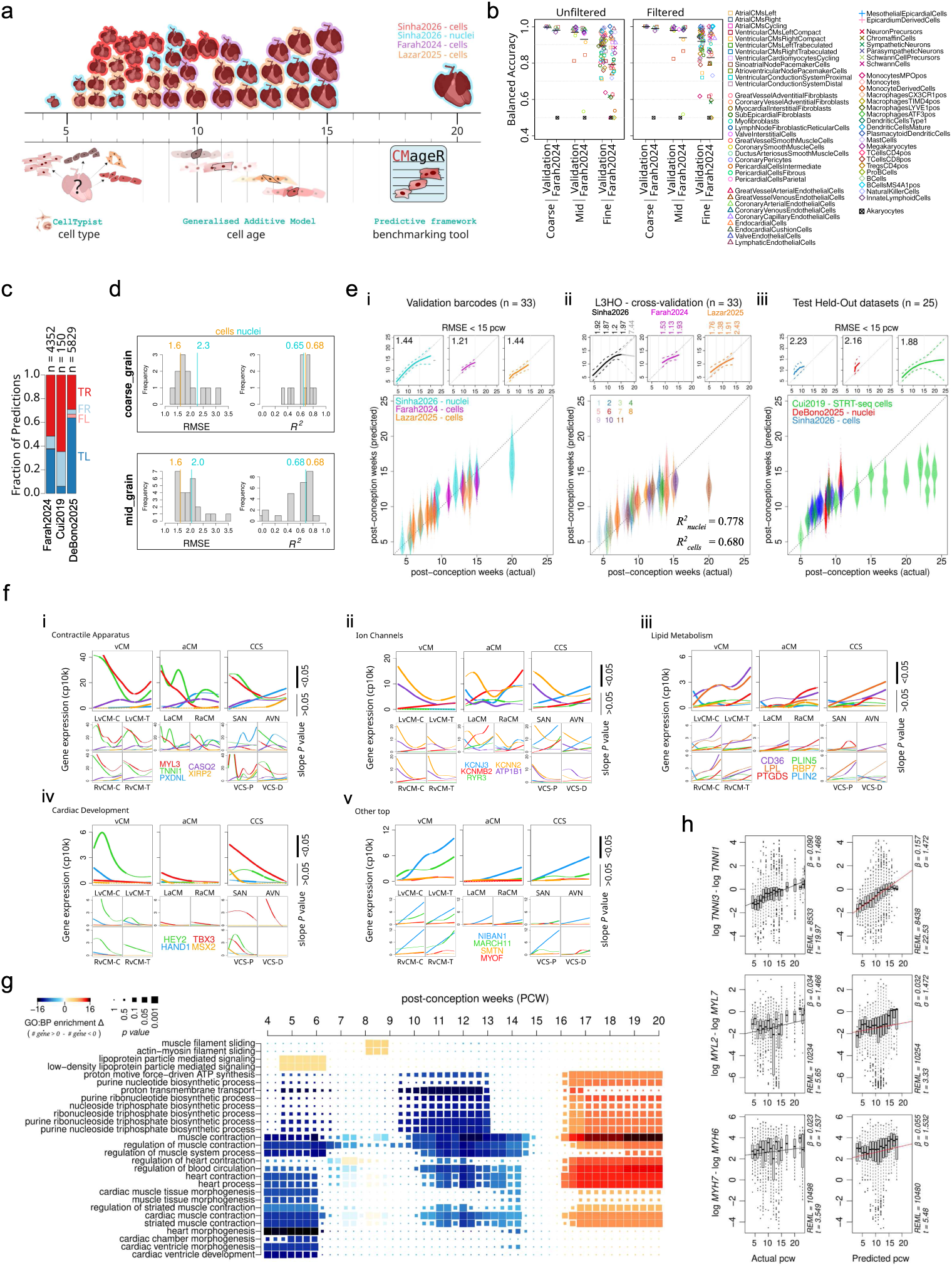
A single-cell transcriptional reference model captures cardiomyocyte maturation dynamics. **A:** Schematic for the two-part computational reference developmental heart model using CellTypist discrete classification, and multivariate generalised additive models (mGAMs) for cell age predictions in the trained model pipieline `CMageR`. The CellTypist models were constructed on a balanced sample of training data from our dataset with a mixture of cells and nuclei. Three discrete classification models were made on the hierarchical granularities of coarse grain, mid grain, and fine grain. In parallel, the mGAMs were constructed on balanced datasets of nuclei from our dataset, and cells from two external datasets (Farah2024[16], and Lazar2025[8]). A developmental age mGAM was constructed for each cell type where data was available, resulting in 6 coarse grain, 13 mid grain, and 55 fine grain models altogether. **B:** Per-class balanced classification accuracies for coarse, mid, and fine grain models in predicting cell types from validation barcodes in our dataset (Sinha2026) and the external held-out scRNA-seq dataset (Farah2024) before and after applying a 20-fold odds-ratio filter. Black horizontal lines show overall model accuracy. **C:** Total population of predicted left or right ventricular cardiomyocytes (vCMs) labelled as True Right (TR), False Right (FR), False Left (FL), or True Left (TL). Red and blue correspond to Right and Left ground truth respectively either from dissected regions (Farah2024, Cui2019[1], or from provided annotations (DeBono2025[39]). Numbers (n) above bars indicate the total sample size of the confidently predicted CMs from fine grain predictions. **D:** Cell age model statistics for the spread of coarse grain and mid grain cell type age predictions shown as histograms of root mean square error (RMSE) and R-squared for samples up to 15 post-conception weeks (PCW) for both sncRNA-seq (nuclei) and scRNA-seq (cells). Medians are shown above each of cells and nuclei. **E:** Distributions of predicted age against actual age (post-conception weeks) in predicted vCMs from fetal data partitions by i, training hearts but unused nuclei and unused cells; ii, leave-3-heart-out (L3H0) out-of-fold (OOF) predictions in train cells; and iii, entirely held out hearts and external data hearts. Predictions were made using the mid grain Ventricular-Cardiomyocytes mGAM where each point is the predicted age of an cell barcode. Subplots above show mean and symmetrical RMSE for individual datasets as solid lines and dashed lines respectively. RMSE values inset are reported for samples < 15 PCW. **F:** Cardiomyocyte-relevant dynamic changes are demonstrated by modelling balloon plots across mid grain, and fine grain cardiomyocyte subtypes showing a selection of nonzero top features associated with cardiomyocyte-relevant functions of i, contractility; ii, ion channels; iii, lipid metabolism; iv, cardiac development; and v, other top markers. The *x* axes cover the 4 to 20 PCW period. Thicker lines denote a significant slope at that period (p < 0.05). **G:** Gene set enrichment results for the top groups of Gene Ontology biological processes (GO:BP) across 51 time points between 4 and 20 PCW illustrating dynamic functional changes within the smoothers generated for the vCM mid grain model features. For each time point, genes with an increasing or decreasing significant change in gradient were enriched for GO:BP using GProfiler2. The color gradient shows the difference between the number of increasing and decreasing genes, while the point size reflects the *p-value*. **H:** Ratios of *TNNI3:TNNI1, MYH7:MYH6*, and *MYL2:MYL7* in vCMs from *in vivo* scRNA-seq were compared with their actual ages, or predicted ages using two competing linear mixed effects models with a modestly improved fit for *TNNI3:TNNI1*, *MYH7:MYH6*, but not *MYL2:MYL7* when modelled by predicted age rather than actual age.

For discrete cell type classification we used the established scRNA-seq logistic regression package CellTypist[29], training 3 independent models on all coarse, mid, and fine grain annotations on a training sample of both cells and nuclei from our dataset only (**Fig. 5b**). Both cells and nuclei were included to train our model for generalisability as performance on cross-platform held-out data was lower when trained on either cells or nuclei alone (**Supp. Fig. S10b**). We applied filtering on the prediction probabilities to improve classification accuracy (see 4), achieving mean per-class balanced accuracies of 1.00, 0.98, and 0.92 on in-house validation cells from the same hearts as training data, and 0.99, 0.94, and 0.83 on coarse grain, mid grain and fine grain cell types respectively on a manually re-annotated external held-out test dataset Farah2024 (**Fig. 5b**, **Supp. Fig. S10d**, **S11**, **Supplementary Table 3**). Even without re-annotation, an accuracy of 0.94 was seen when collapsing our fine grain predictions into the mid-level granularity of the study’s original annotations (**Supp. Fig. S10e**). Furthermore, we demonstrated using dissected-region information as ground-truth that even transcriptionally similar cell types such as L or R vCMs were separable in held-out data with accuracies of between 0.89 and 0.93, and high precisions for RvCMs of between 11 % and 17 % in 10x-scRNA-seq samples (**Fig. 5c**, **Supplementary Table 4**). In predicting fine grain cell types, we also compared our strategy to an alternative scArches label transfer model[30] noting higher accuracies overall (**Supplementary Table 4**, **Supplementary Materials**). Lastly, fine grain accuracy was only marginally affected by the effect of modality or by developmental time (**Supp. Fig. S10f**).

### 2.8 Building a single-cell developmental cardiac clock

We developed our cardiac clock to predict cardiac cell developmental age from singlecell transcriptomics by modelling time (PCW) in each cell type as a response variable to the expressions of different genes as predictors using mGAMs (**Fig. 5a**). For each cell type, an mGAM was trained using gamsel[31] with pre-screened genes binned into trajectory categories to ensure a meaningful feature space across the entire developmental window prior to regularisation (**Supp. Fig. S12a**, see 4). For coarse grain or mid grain, 33 cell types fetal hearts in total from this study and two additional scRNA-seq studies were used and the models were trained using a leave-3-hearts-out (L3HO) cross-validation strategy (**Fig. 5a**, **Supp. Fig. S12c**)[8, 16]. For fine grain cell type models, annotations were not harmonious across all datasets and fine grain mGAMs were only trained on the 11 snRNA-seq hearts from our study, limiting the interpretation of generalisability of the fine grain models.

We assessed model performance by comparing actual PCW to predicted PCW in cells from validation data (training data hearts but unused barcodes), out-of-fold (OOF) predictions from independent L3HO models in training hearts, and from entirely held-out mixed modality hearts from several hearts and studies (**Supp. Fig. S12b**). Under prediction and a marked drop in accuracy was seen at 20 PCW across all cell types with root mean square errors (RMSEs) of between 6.2 and 8.1 weeks, likely resulting from data scarcity as the RMSEs at 20 PCW were halved in validation barcodes for the final model fit when this sample was included (**Supp. Fig. S12b**). For a more meaningful interpretation of our cardiac clock accuracy, we report the performance across `CMageR`’s intended useful temporal interval of below 15 PCW.

Firstly, between 4 and 15 PCW, the models predicted single cell age with an median error of between 1.6 and 2.3 weeks (RMSE) depending on the cell type and model granularity (**Fig. 5d**, **Supp. Fig. S13a**, **Supplementary Table 6**). Secondly, we showed that our models explain an average of 68 % of the variance (*R*^2^) in chronological age through linear regression between actual and predicted age (**Fig. 5d**, **Supplementary Table 6**). A particularly stand-out performance was seen in EpicardialCells using the mid grain model as shown in L3HO OOF predictions in training data (*RMSE = 1.5, R*^2^ *= 0.82*) and in validation nuclei (*RMSE = 1.3, R*^2^ *= 0.89*), agreeing with our previous findings of well-defined epicardial gene expression gradients during development[7]. Lastly, cell age models of very poor performance were Neurons (not Glia), Lymphatic Endothelial Cells, and Leukocytes, likely the result of low cell numbers, unbalanced technical effects, or from underlying heterogeneity within categories during training (**Supplementary Table 6**). We then shuffled mid grain cell labels before re-applying the age-prediction models, the predictive performance dropped substantially (maximum *R*^2^ *< 0.3*) showing cell type specificity of the models (**Supp. Fig. S12e**, **Supp. Fig. S13b**). For a final assessment of the model and comparison against alternative but standard atlas-level integration approaches we constructed a scANVI[32] model on only CMs from our snRNA-seq dataset using scArches[30] (hearts = 11), training a 12-class label transfer model on mid grain vCMs, aCMs, and CCS binned into 4 age groups. In comparison with three gamsel[31] models trained on the same snRNA-seq hearts in vCM, aCM, and CCS clusters (see **Supplementary Materials**), scArches was less accurate across held-out datasets in all CM subtypes (**Supp. Fig. S12d**). While we concede that this alternative approach may be optimised, the results demonstrate that our models’ age-predictive powers are superior to current out-of-the-box integration strategies while avoiding the need for atlas-level integration.

Overall, these findings suggest that within the data-rich first two-thirds of our developmental window, our cardiac clocks predict single-cell age with consistent error and explain a major proportion of chronological age at this time in most cell types with generalisability across sequencing modalities. In addition to use as a predictive tool, the genes selected during model training are themselves biologically informative, since the training process preferentially retains genes that are coupled to developmental progression, so that the model parameters directly reflect the transcriptional programme of CM maturation.

### 2.9 Model parameters reveal functional transcriptional dynamics in cardiomyocyte development

Our model framework can predict the age of an individual vCMs between the ages of 4 and 15 PCW with an RMSE of less than 2.2 PCW in diverse data, resulting in a biological clock that explains at least 68 % of variance due to real chronological age within this time period, depending on the modality (**Fig. 5e**, **Supplementary Table 6**).

To further demonstrate that our models capture true biological signals rather than technical noise, we explored the selected nonzero genes across the four coarse and mid grain CM models, noting many previously described markers of CM maturity and function such as *PDE1C* [33], *LPL*[34], *CASQ2*, *XIRP2* [35], and *TNNI1* (**Fig. 5f**, **Supp. Fig. S12a**, **Supplementary Table 7**). We grouped select top features and known genes associated with CM contraction, conduction, lipid handling, and cardiogenesis, finding that they exhibited different trajectories depending on the CM subtype (**Fig. 5f**). For example, increasing expressions of *KCNJ3* or *PXDNL* in aCM, SAN, and AVN but not in any vCMs, or *RYR3* and *KCNMB2* increasing preferentially in LaCM (**Fig. 5f**). Additionally, the increase in *XIRP2* was more pronounced earlier in RvCMs and VCS-proximal than in other CM types and that vCM age could be described by increasing expressions of *MARCH11*. Our gene trajectories also show that lipid handling is an important predictor of ventricular, but not atrial CM developmental age as noted in *LPL* and *CD36* expression, while the decreasing slopes of developmental transcription factors *HAND1*, *HEY2*, *MSX1*, and *TBX3* were varied between the different chambers or CM subtypes (**Fig. 5f**). Interestingly, *TNNI1* expression - often a surrogate for CM immaturity, tends towards zero in working aCMs but not in working vCMs (**Fig. 5f**). Lastly, ranked by cross-validation importance, the top 5 markers overall were *MB*, *MYL3*, *NIBAN1*, *RBP7*, *PTGDS*. As a top positive predictor of CM developmental age, *NIBAN1* is particularly interesting: CMs have the highest expression of NIBAN1 among all cell types documented in the Human Protein Atlas[36], and it is one of the most abundant proteins expressed in CMs[37] and yet research on its function in CM maturation and ageing is wholly absent. As it has a key role mediating pro-survival responses to cell stress[38] and apoptosis, it is a prime candidate for further research in CM growth and maturation.

Providing further evidence of clear biological relevance in our entire selected feature space, a dynamic over-representation analysis (ORA) using the significant derivatives of the gene slopes was carried out for the features of VentricularCardiomyocytes mGAM giving a meaningful representation of the dynamic changes in vCM focus across the developmental window (**Fig. 5g**, see 4). Following 4 PCW, the vCMs are downregulating a large number of genes that are involved in heart and ventricle morphogenesis while upregulating genes associated with processes of lipoprotein signalling. This is coupled to a more drawn-out decrease in processes of purine nucleotide biosynthesis or proton transmembrane transport (including genes *ENO1*, *ATP1B1*). Unsurprisingly, this analysis also revealed a shifting landscape of cardiac and muscle-related programs as genes associated with muscle contraction related processes were highly dynamic throughout (**Fig. 5g**). Towards the end of the window, a strong enrichment of regulation of cardiac muscle contraction and other contraction terms was seen. In summary, the period covered here shortly after looping and formation of the four chambers is reflected in this continuum as loss of key developmental programs involved in gross morphology, the establishment of a metabolic microenvironment to sustain continuous cardiac contraction, and the dynamic rearrangement of components involved in contractile apparatus. This progression captures a critical and previously unresolved phase of human cardiac development at molecular resolution.

Finally, we established that our vCM model uncovers biological age at a single cell level, describing the heterogeneity of cell ages within samples by showing that predicted ages compared rather than actual sample age was a better predictor for known metrics of CM maturation in single cells. This was demonstrated in vCMs from all datasets by modelling the ratios of *TNNI3:TNNI1*, *MYL2:MYL7*, and *MYH7:MYH6* using two competing linear mixed-effects models in response to either predicted or actual age, modelling the intercepts individually within 10x-snRNA-seq, 10x-scRNA-seq, and STRT-seq data (**Fig. 5h**). While both predicted age or actual age captured a significant positive increase in all ratios, our model’s output exhibited modestly better predictive power and improved overall model fit when compared to fixed chronological time in both ratios of TNNI, and MYH, but not MYL (**Fig. 5h**). Specifically, the relationship between *TNNI3:TNNI1* and chronological time was shallower with a poorer model fit (β = 0.090 ±0.006, REML = 8533, t = 19.97) than that observed between *TNNI3:TNNI1* and our predicted biological time (β = 0.157 ±0.007, REML = 8438, t = 22.53). This improvement was seen despite the already-poor association between the *TNNI* ratio and chronological time (*R*^2^ < 0.14) (**Supp, Fig. S13c**). Interestingly, *TNNI1* was not a selected feature in the mid grain vCM model and its independent dynamics here substantiates our clock’s alignment of biological age. The discrepancy in MYL may be explained by cross-modality differences and data scarcity as the STRT-seq dataset Cui2019[1] showed an increasing trend in the *MYL2:MYL7* ratio particularly in later time points, whereas our training data vCMs exhibited mixed trends causing both *MYL2* and *MYL7* to be omitted from the vCM and CM models (**Supp. Fig. S13c**). Lastly, we note that the *TNNI3:TNNI1* ratio plateaus after 15 PCW, representing a possible biological phenomenon in vCM maturation, and offering a partial explanation for poor predictive performance after this age (**Supp, Fig. S13c**).

Together, these findings suggest that our developmental cardiac clock captures the continuum of biological age in CMs between 4 and 15 PCW, providing a more robust signal to noise ratio than currently used ratiometric measures of contractile apparatus, or even by their sampled chronological age. Furthermore, the functional assessment of the selected genes demonstrates that our model covers a critical window of cardiac cell type specification and maturation following the formation of the 4 chambers. By predicting cell type followed by cell age, we can place unseen cells along a meaningful continuum of cardiac development.

### 2.10 Benchmarking purity, diversity, and maturity in hPSC-CMs

We used the `CMageR` pipeline to benchmark hPSC-CM models by gathering published scRNA-seq data from 10 peer-reviewed articles, spanning 83 conditions with varied differentiation protocols of putative CM subtypes and maturities[40–49] (**Supplementary Table 8**). Starting a low resolution, we found some coarse grain CM predictions in *in vitro* cells at early time points. As it is unlikely that true CM-like cells exist in culture immediately after day 0, we calibrated a heuristic confidence threshold of 0.2 probability using day 3 and day 4, aligning cardiac induction timing in many hPSC-CM differentiation protocols (**Supp. Fig. S14b**). The median probability of all in vitro conditions after day 10 was higher than this 0.2 threshold, except a commercial 2D protocol by Cellapy (Ou2021)[42], suggesting that this protocol may take longer to generate high-fidelity CMs (**Supp. Fig. S14b**). Above this threshold, our model revealed a wide spread in CM purity that was described by a significant positive monotonic relationship with culture day across all *in vitro* conditions (*Spearman’s ρ = 0.71, p < 0.001, n = 83*) (**Fig. 6a**). One of the faster protocols to achieve a high purity was the embryoid body bioreactor protocol using the H9 cell line (Selvakumar2024)[44]. Notable exceptions to the positive trend were organoids or complex 3D constructs. For example, early organoid models by (Schmidt2023)[49] were of high CM purity after only 7.5 or 9.5 days of culture time, comparable to that of monoculture protocols with low-glucose selection steps such as Funakoshi2021[40] at days 20 and 32, or Galdos2023[41] from day 30 (**Fig. 6a**). Unsurprisingly, assembled EHTs in Cheng2023[46] or microtissues (MT1 and MT2) in Giacomelli2020[45] with long culture times were of lower CM purity, both requiring complex protocols or assembled from a mixture of cell types. This was particularly low in the vCM EHTs by (Cheng2023)[46] with a mean cardiomyocyte purity as low as 12 % ±7 % (mean, sd, *n = 3*) in their Cardiac Organoid Chambers (COC) (**Fig. 6a**).

**Fig. 6:**
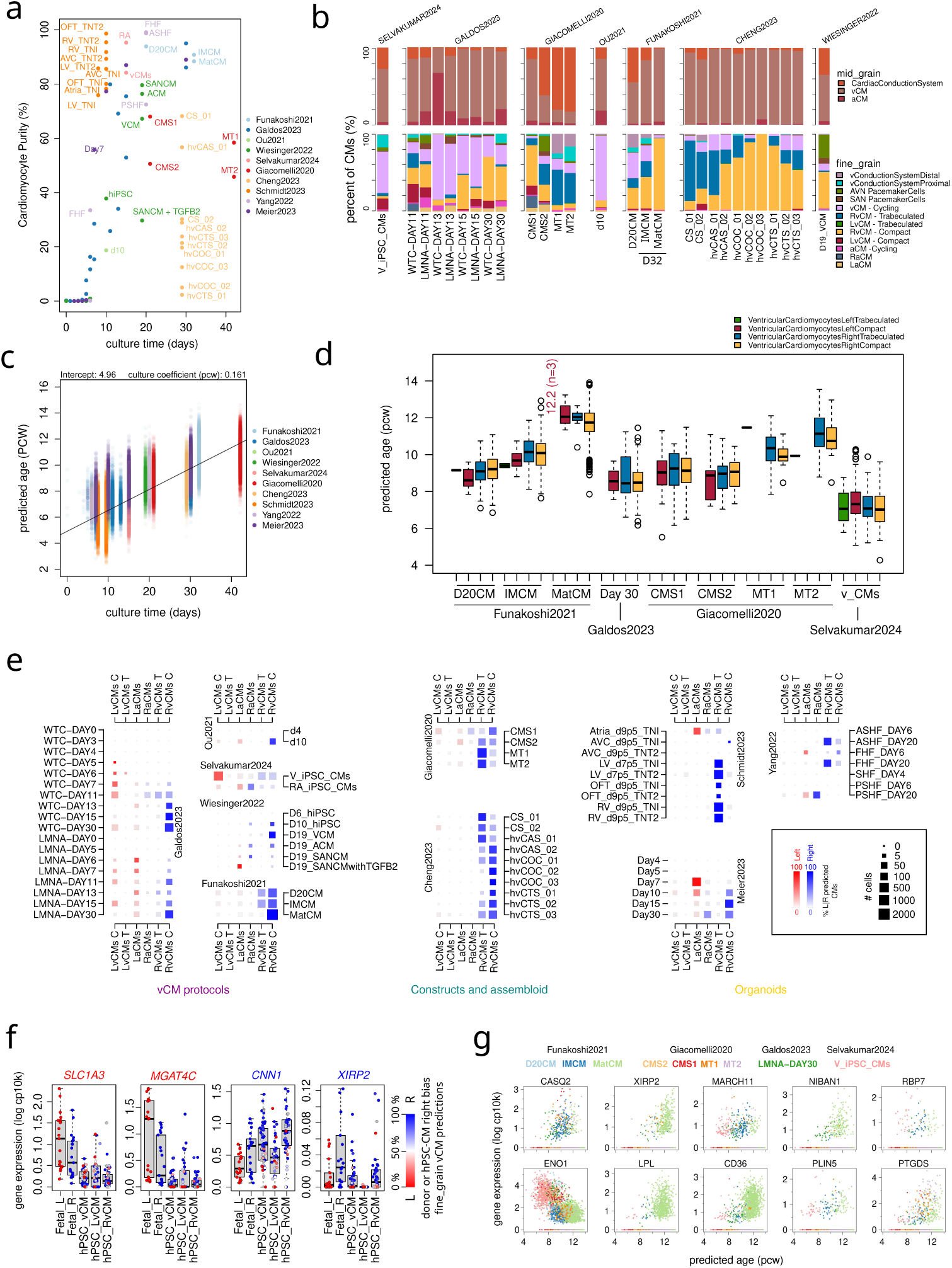
Predicting cell type and cell age of hiPSC models reveals a right transcriptional bias in ventricular cardiomyocytes. **A:** The fraction of cardiomyocytes (CMs) predicted in iPSC-Cardiomyocyte models from different differentiation days across different protocols and aims. There was a significant positive monotonicc relationship between cardiomyocyte purity and culture day (*Spearman’s ρ = 0.71, p < 0.001, n = 83*) **B:** CM subpopulation plots showing predicted cell type composition of mid grain and fine grain annotations across multiple datasets from protocols that report ventricular phenotypes. Populations are represented as a fraction of all high-confidence CM predictions (coarse grain CM prediction probability *>* 0.2). **C:** The association between predicted developmental time and culture time in days of CMs extracted from all protocols and conditions. A linear model was fitted on all CMs from the hPSC-CM model datasets using predicted age as a dependent variable, and culture time as independent variable generating the two coefficients of intercept and culture time. Significant correlation between predicted age of CMs and culture time was seen (*Pearson’s r = 0.77, P < 0.001, n = 143,270*). **D:** Box and whisker plot showing the predicted ages of left (L) and right (R) ventricular cardiomyocyte (vCM) subtypes within 4 datasets and selected conditions. **E:** L vs R balance of vCMs and atrial cardiomyocytes (aCMs) across all *in vitro* datasets and conditions (n = 83). Balance was calculated as the fraction of each cell type within all L or R labelled vCMs. Our representation shows both balance and total numbers. Conditions where no CMs were found were omitted. **F:** Mean expressions of selected subset of LvCM genes *SLC1A3* and *MGAT4C*, and *RvCM* genes *CNN1* and *XIRP2* plotted across the LvCM and RvCM clusters from individual fetal donors or the 83 different hPSC-CM conditions either as all vCM predicted cells, or separately as LvCM or RvCM cells. **G:** Selected top features from the vCM age model are associated with predicted developmental age of the hPSC-CMs. **Study condition glossary**: **Selvakumar2024** - V iPSC CMs (Ventricular induced pluripotent stem cell cardiomyocytes), RA iPSC CMs (Retinoic Acid+ induced pluripotent stem cell cardiomyocytes); **Galdos2023** - WTC (WTC cell line), LMNA (SCVI-111 cell line); **Giacomelli2020** - CMS* (cardiomyocyte only replicate), MT* (microtissue replicate); Ou2021, d0, d2, d4, d10 (days 0, 2, 4, & 10); **Funakoshi2021** - D20CM (day 20 cardiomyocytes), IMCM (day 32 - maturation media), MatCM (day 32 + maturation media); **Cheng2023** - CS (cardiac spheroids replicate), hvCAS (human ventricular cardiac anisotropic sheet replicate), hvCOC (human ventricular cardiac organoid chamber replicate), hvCTS (human ventricular cardiac tissue strip replicate); Wiesinger2022 - d0, d4, d5, d6, d10, d19 (days 0, 4, 5, 6, 10 & 19), VCM (day 19 ventricular cardiomyocytes), ACM (day 19 atrial cardiomyocytes), SANCM (day 19 sinoatrialnode cardiomyocytes), SANCM+TGFB2 (day 19 sinoatrialnode-transitional zone cardiomyocytes); **Schmidt2023** - d7p5 & d9p5 (days 7.5 & 9.5), AVC (atrioventricular canal), LV (left ventricle), RV (right ventricle), OFT (outflow tract); **Yang2022** - FHF(first heart field), SHF (secondary heart field), ASHF (anterior secondary heart field), PSHF (posterior secondary heart field)

In assessing the mid grain CM age model outputs, we found a significant linear relationship between culture time and predicted age (**Fig. 6c**) (*Pearson’s r = 0.77, p < 0.001, n = 143,270*). By fitting a simple linear model to predicted cell ages, a coefficient of 0.161 PCW suggests that the rate of hPSC-CMs maturation and growth parallels real development, at approximately 1.1 days per day of culture (**Fig. 6c**). Unsurprisingly, a weaker positive correlation with predicted age was also seen with the co-correlating variable of CM classification probability (*Pearson’s r = 0.13, p < 0.001, n = 143,270*). Broadly, the results agree with author descriptions of the stem-cell models as the lowest predicted ages were seen in early-developmental multi-chamber organoid models[49], while the oldest predicted samples were associated with conditions with specific aims of maturation[40] (**Fig. 6c**). The alignment of our model’s output with published accounts of hPSC-CMs provides evidence of translation and generalisation into *in vitro* systems, offering an unbiased biological signal of developmental age or maturation.

### 2.11 In vitro vCMs are transcriptionally right-chamber biased

We then explored the CM subtypes within the datasets that were reported to contain vCMs[40–45]. On the whole, agreeing with the claims of the authors, the most abundant mid grain cell type predicted in these datasets was vCMs (**Fig. 6b**). However, aCM predictions were often seen at earlier stages of differentiation (**Supp. Fig. S14c**), and a large proportion of CCS cells was seen in some cases such as in Giacomelli2020 microtissue (MT) replicates (**Fig. 6b**), or making up the majority of the multichamber organoids[49] at 7.5 and 9.5 (**Supp. Fig. S14c**). Curiously, the Giacomelli2020 CCS predicted cells were resolved as trabeculated vCMs using the fine grain model, suggesting some transcriptional similarity and confusion between the two classes in vitro (**Fig. 6b**, **Supp. Fig. S11**), while the organoids in Schmidt2023 may represent very early cardiogenic phases resulting in out-of-domain misclassification. However, given their published characterisations, we expected to see greater differences between the chamber-specific models.

Further detail was revealed using the fine grain classification results, focusing on several vCM studies that reported increased maturation or compaction in hPSC-CMs (**Fig. 6b**). Funakoshi et al., (2021)[40] showed increased CM compaction as a result of switching metabolism to fatty acid oxidation and our predictions agree, revealing a switch between trabeculated-like vCMs and compact vCMs (**Fig. 6b**). Giacomelli et al., (2020)[45] reported an improved maturation in the assembled MT constructs when compared to CMs alone (CMS1 and CMS2). However, unlike the Funakoshi2021 protocol, our predictions suggest that this resulted not in a change in vCM cell type but of cell age, increasing the developmental age in vCMs in the construct by 1-2 weeks (**Fig. 6d**). Broadly, we saw that VentricularCardiomyocytesCycling were particularly abundant across earlier timepoints as a large population of most hPSC-CM models. Interestingly, despite the low CM purity within 3D constructs from the Cheng2023 dataset, their CM subpopulations were almost entirely ventricular, with many compact vCMs predicted in the COC and cardiac tissue strip (CTS) condition[46]. Lastly, we saw broader evidence of retinoic acid (RA) driving increased atrial and pacemaker cell specification in the protocols Selvakumar2024 and Wiesinger2022, aligning with current understanding in the field (**Supp. Fig. S14c**).

Within these vCMs we found that the predicted developmental age was highest with a mean of 12.0 PCW in the mature CM (MatCM) condition from the Funakoshi2021 dataset, followed by vCMs from the Giacomelli2020 data (10.7 PCW) when using the mid grain VentricularCardiomyocyte model (**Fig. 6d**). By partitioning these results by fine grain, we found that a handful of compact LvCM predicted cells in the Funakoshi2021 MatCM condition were the oldest, with a mean developmental age of 12.2 PCW (n cells = 3), placing them 2.5 weeks older than vCMs in the immature CM (IMCM) condition cultured for the same period of 32 days, but lacking maturation media (**Fig. 6d**). Interestingly, despite the high purity and long culture time, the CMs from Galdos2023 at day 30 remained developmentally young, with a mean age of 8.5 PCW. Even younger still at 7 PCW were the vCMs in Selvakumar2024 15-day VCM condition (**Fig. 6d**). This suggests that while there is an association between chronological culture time and CM transcriptional age, it can be disconnected and accelerated through culture conditions.

Finally, while we highlighted populations of trabeculated or compact vCMs in many protocols, we calculated a L / R ratio for both atrial and ventricular CMs and we found that the majority of vCM predictions were right-chambered, even in the later time points of TBX5+ lineage cells where LvCMs were reported as the primary population[41] (**Fig. 6e**). The major exceptions were the VCM condition in Selvakumar2024 data, and to a lesser extent, the earlier or mid time points in Galdos2023 data and in the epicardioid Meier2023[47] data (**Fig. 6e**). Interestingly, this bias did not appear to affect atrial predictions. We showed that this was not a technical artefact from this model as similar results were seen using 3 additional independent atlas-level integration approaches using scArches[30] trained on 1, all fine grain annotations for holistic classification (k = 63); 2, all CMs at fine grain to reduce noise (k = 12); or 3, CMs grouped into LvCM or RvCM or “other CM” within every donor to allow for confusion between LvCM and RvCM between different time points (k = 36) (**Supp. Fig S15**). To better understand these findings, we compared the expressions of the top L and R genes in our in vivo donors with predicted vCMs across the 83 conditions in the hPSC-CM models and found that stem cell models highly express RvCM genes such as *CNN1*, or *XIRP2*, but have lower expression of LvCM genes like *SLC1A3* or *MGAT4C*, with distributions spread by the LR bias of the overall hPSC-CM culture (**Fig. 6f**, **Supp. Fig. S15**). We were able to see that LvCM-predicted hPSC-CMs were typically lower in R gene expressions and higher in L gene expressions but with values falling short still of true LvCMs *in vivo*, validating the model outputs (**Fig. 6f**, **Supp. Fig. S15**). Lastly, we found that many of the top predictor genes of vCM age were upregulated specifically in hPSC-CMs predicted to be older (**Fig. 6g**). In particular, the increase in *LPL* and *CD36* and a reduction in *ENO1* with age in Funakoshi2021 MatCM condition is evidence that the conditioning media applied increased fatty acid oxidation as expected where we can now tie this shift to a particular developmental stage. Simultaneously, we see increases in *NIBAN1* with predicted age of stem cell models suggesting that it is meaningful as a marker of maturity *in vitro* as well as *in vivo* (**Fig. 6g**).

As a proof-of-concept, we have shown that our models capture both the expected shifts in hPSC-CM populations as described by authors, and the expected changes in developmental age in association with both culture time and well-defined maturation protocols. However, the predictions from our reference pipeline `CMageR` also revealed disconnects with reported characterisations, including models with unintended subpopulations, low maturities, or transcriptional right-chambered bias.

## 3 Discussion

In this study, and in our sister study we have combined single-cell with spatial transcriptomics[9] to present the most temporally resolved single-cell atlas of the human fetal heart and great vessels to date through the first and second trimesters. Cell type definitions were based on anatomical location and functional identity along with computational cluster boundaries rather than computational predictions alone, with better clarity compared to previous attempts[1–7, 50]. Beyond cataloguing cellular diversity at unprecedented scale, we demonstrate that this atlas can serve as an active reference framework: enabling transcriptional benchmarking of the rapidly expanding repertoire of cardiac stem cell models, from organoids and engineered heart tissues to assembloids, at both the level of cell identity and developmental maturity. While we focused on our vCM model, it is pertinent to point out that our models were trained for all cell types with acceptable predictive performance in many, including endothelial cells, epicardial cells, or even glial cells, each with specific developmental trajectories waiting to be assessed. Even in the well-studied trajectory of developmental age in vCMs we noted several novel markers of vCM maturity, suggesting great value in exploring the other less-studied cell types presented here.

### 3.1 Left and right chamber cardiomyocyte identity

We presented findings on the distinct characteristics of right-left chamber identities for CMs during development with implications for stem cell models and cardiac regeneration. Firstly, an atrial right-left signature emerged more readily distinguishable compared to the ventricular left-right signature. This observation is intriguing, given the aCMs are thought to share the same precursor pool of cells, yet they show a clearer distinction compared to LvCMs and RvCMs, which are believed to develop from different progenitors in first and second heart fields. We explored the role of *PRRX1*, a marker that distinguishes between the ventricular chambers and has previously been suggested to inform the right-left axis during heart looping[51]. However, its role in RvCM identity is not clear. Here we have shown that its knockdown in hPSC-CMs results in a reduction in some key RvCM genes but further studies are required to conclude the extent of its role in chamber specification. Interestingly, our study also found that many RvCM genes including *PRRX1* were not markers for RvCMs in the adult human heart[18], implying differential roles specifically for both these genes and CM function in development. This contrasts with our understanding of physiological and mechanical differences between the ventricles in adults, such as the LV’s higher pressure and thicker walls compared with the RV. To explain this, their similar transcriptional profiles may be attributed to balanced mechanical parameters as both ventricles experience comparable end-diastolic wall stress and wall stress due to compensatory variations in wall thickness, ensuring uniform activation of mechanosensitive pathways. Additionally, similar sarcomere lengths (1.8–2.2 µm) help standardize stretch-sensitive signalling and transcriptional responses. Despite apparent developmental differences, adult uniformity in gene expression may impart resilience against region-specific pathologies and could partially explain why both ventricles exhibit similar molecular changes in advanced stages of certain cardiac diseases. Secondly, as hPSC-CMs are evidently developmental, our findings on right-chambered bias in hPSC-CM vCMs may impact how we select appropriate models for drug discovery, and disease modelling particularly in congenital heart defects. Genetic conditions may be chamber-specific such as arrhythmogenic right ventricular cardiomyopathy, or Brugada syndrome that affect the right ventricle, or even heart failure and ischaemia following myocardial infarction (MI) that predominantly affect the left ventricle. These findings are also highly relevant for cell-based therapies as while a convergent adult identity suggests that environmental pressures may eventually drive vCMs towards a single adult molecular profile, it is unknown whether matching LvCM or RvCM with host tissue ventricles is necessary for efficacy or cell retention of grafts following MI.

### 3.2 Right bias in stem cell derived ventricular cardiomyocytes

Transcriptional right bias and RvCM predictions made here on hPSC-CM data contradicts with current developmental understanding, particularly when observed in the TBX5+ hPSC-CMs[41]. Clear evidence using lineage tracing has shown that TBX5+ CMs from the first heart field (FHF) contribute to the left ventricle and atria, but not the right ventricle in mice[52]. We offer two possible explanations for these predictions that warrant further study. Firstly, TBX5 may not preclude RvCM differentiation in vitro. Transcriptionally, TBX5 appears to be present albeit at lower expression in the human right ventricle in MERFISH data (Farah et al. 2024), (**Supp. Fig. S5b**), and in RvCMs seen in scRNA-seq from data presented here and in previous studies[1, 16], as well as in murine RvCMs[53] (**Supp. Fig. S15e**). From this evidence it is clear that *TBX5* mRNA is not specific to mammalian left-chamber vCMs *in vivo*, suggesting that LvCM-specific activity at the protein-level may be driven by dosage differences or controlled through post-transcriptional regulation. Indeed, research has outlined two miRNAs (miRNA10a / b) that bind specifically to the 3’ untranslated region of TBX5, reducing the protein but not mRNA expression[54]. Further studies need to be carried out to determine whether *in vitro* systems share any post-transcriptional mechanisms, or whether any reporter constructs used in hPSC-CM studies may interfere with them. Secondly, even if *TBX5* + cells are initially and irreversibly LvCM-fated, the conditions imposed on cells during culture may still promote a right-like transcriptional identity through paradevelopmental pathways or resulting from the absence of the in vivo LV niche. Well-established culture conditions used by many laboratories may drive the transcriptional right-bias evident at many later time points in culture regardless of putative specification. Lastly, it is possible that some predictions are simply incorrect. However, we intentionally validated our model on equivalent held-out scRNA-seq data across different chambers, showing that only 17 % of RvCM predictions were actually vCMs from the LV (**Fig. 5c**).

### 3.3 Limitations of the developmental cardiac clock

While we have shown the usefulness of the model output in its current state, there are several limitations and considerations. Firstly, it is solely parametrised using scRNA-seq data and although necessary for dissecting heterogeneous cultures, scRNA-seq data is often noisy and much more expensive than bulk RNA-seq. Secondly, as it is trained on only transcriptomics, the model is missing key epigenetic predictors as seen in established epigenetic clocks. Thirdly, the parametrisation and structure of the current model likely needs improving as performance is poor in extrapolation, particularly at and beyond the upper boundary of our assessed developmental window. That being said, we find that even the most mature hPSC-CM model assessed here at 12 PCW was within our model’s well-captured predictive period and considerable improvements may still be made in cultures before stem cell models approach the 15 PCW mark. Until then we argue that the predictions are still useful. There are many known, latent or technical factors that may contribute to the error and output of the age models. Between 4 and 15 PCW, the model RMSEs were relatively consistent, often under or around 2 weeks. While not small, a major component of this can be attributed to inaccuracies in clinical estimation, collection, and reporting of fetal ages in gestational age (weeks since the first day of last menstrual period) or post-conception weeks (GA - 2 weeks). With a variance in ovulation day estimates by 5-6 days, misreported or rounded values between collection and research teams, and the potential for overestimation of the whole sample by 14 days, clinical estimates are inherently erroneous. In the context of high baseline error, the median RSME of 2 weeks reported here between 4 to 15 PCW is reasonable, especially for single-cell age prediction. While Carnegie staging may also be used it does not cover the period after 8 PCW. The within-sample distributions of age are also likely a combination of technical error from scRNA-seq data, and biological error as a result of maturation or growth heterogeneity within each heart. We provided some evidence that our model captures biological error by providing a modestly better explanation than PCW for the spread of maturation-associated switches in contractile apparatus in vCMs.

Beyond 15 PCW our temporal model approached a predictive plateau despite the steps taken to limit technical biases and select features across the entire developmental window. While the simplest explanation for this is in the imbalance of data, namely scarcity of data at the later timepoints, there may be alternative reasons for this phenomenon. It may be a biological upper limit of maturation as genes cannot increase in expression indefinitely, and cells will be transcriptionally fully mature at some point in time, for example, as seen in the flattening of the *TNNI3:TNNI1* ratio in vCMs. Predicting the age of cells beyond this point may only be possible by including postnatal data to model the additional time-dependent processes of growth or ageing. This limit may also indicate insufficiency in the feature space, the transcriptomics-only approach, or the model framework itself. We briefly reviewed age predictions in adult data and noted that the 15 to 20 PCW plateau persisted in adult predictions while the intercept of the linear model for culture time in hPSC-CMs was at the lower limit of our dataset age, and not more towards the expected value of 0. These observations suggest boundary effects, either biological or technical, that may be improved through alternative model frameworks. Further comparisons need to be made against multiple linear regression or support vector regression models that may improve extrapolative capacity beyond the current mGAM structure. Nonetheless, despite these caveats, our model presented here as a proof-of-concept behaves consistently across the modalities assessed and both in vivo and in vitro data even to the point that our model revealed mislabelled samples deposited on GEO by predicting the order of the ages correctly. This was confirmed after communication with the authors[39], where they concluded this to be the case through orthogonal assessment of *XIST* expression from the barcodes. This demonstrates a real utility not only in benchmarking stem cell models, but also as part of a standard data curation pipeline. Future iterations of this model should include balanced data well into adulthood, comparisons of additional model structures, additional modalities including spatial transcriptomics, and an optimised feature space.

### 3.4 Conclusion

Many researchers working with hPSC-CMs are actively using, assessing, and improving their models but characterisations are often fragmented or study-specific. As a proof-of-concept to unify comparisons across the field, we sought to develop a transcriptional ground-truth model of cell type, and biological age for cardiac cells in a key developmental window. Through a preliminary benchmark of many hPSC-CM models, we demonstrated that this allowed for an unbiased dissection of the cell type composition within cultures, and with a single clear real-world metric of developmental age, has addressed a long-standing question of stem cell and developmental biologists: how old are our cardiac stem cell models? While hPSC-CM maturity and type may continue to be assessed through functional and morphological characteristics and supervised analysis of individual components, we hope that `CMageR` and its future versions may be used as standard reference, improving our ability to benchmark cardiac stem cell models, especially within the bustling era of new approach methodologies (NAMs). Finally, while we have utilised our atlas by targeting the field of stem cell models, the resource presented here may also serve as a reference for differences between human and animal models. For example, we noted that several markers were used to mark the entire CCS in mice[17, 55]. However, within the developing human heart, no marker was identified that could capture all the components of CCS in a sensitive and a specific manner. Another difference observed during the analysis was a lower expression of *TCF21* in human epicardial cells, contrasting with well known expression in zebrafish where *tcf21* is used as an epicardial lineage marker, complicating translational regenerative research from zebrafish studies[56]. As noted throughout, many opportunities now exist to tackle technical limitations encountered in our study, notably the application of single-cell resolution spatial transcriptomics methods. Until then and despite these challenges, the insights obtained from this atlas are anticipated to have broad applications in cardiovascular research and contribute to a deeper understanding of human heart development.

## 4 Methods

### 4.1 Ethics and human embryonic and foetal heart tissues

Ethics The embryonic and foetal heart samples corresponding to the donor IDs BRC2251, BRC2253, BRC2256, BRC2260, BRC2262, BRC2263, C82, C83, C85, C86, C87, C92, C94, C97, C98, C99, and C104 were provided from terminations of pregnancy from Cambridge University Hospitals NHS Foundation Trust under permission from NHS Research Ethical Committee (96/085). The donor IDs Hst33, Hst36, Hst39, Hst40, Hst41, Hst42, Hst44, and Hst45 were provided from the MRC/Wellcome Trust Human Developmental Biology Resource (University College London (UCL) site REC reference: 18/LO/0822; www.hdbr.org). Sample names and that information are listed in Supplementary Table 1.

### 4.2 Sample collection and processing

The processing of samples with donor IDs of BRC2251,BRC2253, BRC2256, BRC2260, BRC2262 and BRC2263 were described previously[7]. Briefly, these samples were stored at 4 °C overnight in Hibernate-E medium (ThermoFisher Scientific). The next day, the apex and the base of the heart were dissected and separately dissociated using 6.6 mg/mL Bacillus licheniformis protease (Merck), 5 mM CaCl2 (Merck), and 20 U/mL DNase I (NEB). The mixture was triturated on ice for 20 seconds every 5 minutes until clumps of tissue were no longer visible. The digestion was stopped with ice-cold 10% fetal bovine serum (FBS, ThermoFisher Scientific) in phosphate-buffered saline (PBS, ThermoFisher Scientific). Cells were then washed with 10% FBS, resuspended in 1 mL PBS and viability assessed using Trypan blue. Cells were submitted for 10x library preparation with v3.0 chemistry for 3’ single-cell sequencing on a NovaSeq 6000 (Illumina) at the Cancer Research UK Cambridge Institute.

For the processing of donors corresponding to the donor IDs C86, C94, C97 and C99, samples were stored at 4 °C overnight in HypoThermosol preservation solution (Merck). Tissue was first minced in a tissue culture dish using scalpel. Minced tissue was digested with type IV collagenase (final concentration of 3 mg/mL; Worthington) in RPMI (Sigma-Aldrich) supplemented with 10% fetal bovine serum (FBS; Gibco), at 37°C for 30 min with intermittent agitation. Digested tissue was then passed through a 100-µm cell strainer, and cells were pelleted by centrifugation at 500g for 5 min at 4°C. Cells were then resuspended in 5 ml of red blood cell lysis buffer (eBioscience) and left for 5 min at room temperature. It was then topped up with a flow buffer (PBS containing 2% (v/v) FBS and 2 mM EDTA) to 45 ml and pelleted by centrifugation at 500g for 5 min at 4°C. The resuspended cell solution was filtered through a 70-*µ*m cell strainer (Corning), and live cells were manually counted by Trypan blue exclusion. Dissociated cells were first incubated with 5 *µ*L of FcR blocker for 5 min at room temperature and stained with anti-CD45 antibody (BV785 anti-human CD45 antibody, BioLegend, 304048) and DAPI (Sigma-Aldrich, D9542) prior to sorting. DAPI was used at a final concentration of 2.8 *µ*M, and all antibody solutions were used at a final concentration of 5 µl per 100 *µ*l cell suspensions containing fewer than 5 million cells. DAPICD45+ and DAPI-CD45-populations were sorted by FACS using MA900 Multi-Application Cell Sorter (Sony) and its proprietary software (Cell Sorter Software v3.1.1). Sorted cells were loaded on the Chromium Controller (10x Genomics) with a targeted cell recovery of 5,000–10,000 per reaction. Single-cell cDNA synthesis, amplification, gene expression library was generated according to the manufacturer’s instructions of the Chromium Next GEM Single Cell 5’ Kit v2 (10x Genomics). Libraries were sequenced using NovaSeq 6000 (Illumina) at Wellcome Sanger Institute with a minimum depth of 20,000–30,000 read pairs per cell.

Samples used for single nuclei isolation were flash-frozen (unembedded) or frozen in OCT and stored at -80 °C, or formalin-fixed and subsequently embedded in paraffin blocks. All tissues were stored and transported on ice at all times until freezing or tissue dissociation to minimise any transcriptional degradation. Single nuclei were obtained from flash-frozen tissues using sectioning and mechanical homogenization as previously described. 5-10 mm thickness frozen tissues were first sectioned with cryostat in a 50 µm thickness section. All sections from each sample were homogenised using a 7 ml glass Dounce tissue grinder set (Merck) with 8–10 strokes of a tight pestle (B) in homogenization buffer (250 mM sucrose, 25 mM KCl, 5 mM MgCl2, 10 mM Tris-HCl, 1 mM dithiothreitol (DTT), 1 × protease inhibitor, 0.4 U *µ*l-1 RNaseIn, 0.2 U *µ*l-1 SUPERaseIn, 0.05% Triton X-100 in nuclease-free water). Homogenate was filtered through a 40-µm cell strainer (Corning). After centrifugation (500g, 5 min, 4 °C) the supernatant was removed and the pellet was resuspended in storage buffer (1 × PBS, 4 % bovine serum albumin (BSA), 0.2U *µl*^−1^ Protector RNaseIn). Nuclei were stained with 7-AAD Viability Staining Solution (BioLegend) and filtered through 20-*µ*m cell strainer (CellTrics). Positive single nuclei were purified by fluorescent activated cell sorting (FACS) using MA900 Multi-Application Cell Sorter (Sony) and its proprietary software (Cell Sorter Software v3.1.1). Nuclei purification and integrity were verified under a microscope, and nuclei were manually counted by Trypan blue exclusion. Nuclei suspension was adjusted to 1000-3,000 nuclei per microlitre and loaded on the Chromium Controller (10x Genomics) with a targeted nuclei recovery of 5,000-10,000 per reaction. 3’ gene expression libraries. Libraries were sequenced using NovaSeq 6000 (Illumina) at Wellcome Sanger Institute with a minimum depth of 20,000-30,000 read pairs per nucleus.

### 4.3 Read mapping

After sequencing, samples were demultiplexed and stored as CRAM files. Each sample of single-cell data was mapped to the human reference genome (GRCh38-2020-A) provided by 10x Genomics using the CellRanger software (CellRanger v.6.0.2 or v.6.1.0, or CellRanger ARC v.2.0.0) with default parameters. Each sample of Visium was mapped to the human reference genome (OCT: GRCh38-3.0.0, FFPE: GRCh38-2020-A) the SpaceRanger software (OCT: v.1.1.0, FFPE: v.2.0.0, Visium HD: v.3.0.1) with default parameters. For Visium samples, SpaceRanger was also used to align paired histology images with mRNA capture spot positions in the Visium slides. Part of the single-cell samples were mixed with different donors after the nuclei isolation for cost-efficient experimental design (Supplementary Table 1) and computationally demultiplexed using Soupercell[57] (v.2.0) based on genetic variation between the donors.

### 4.4 Preprocessing and quality control

For the single-cell transcriptome data, the CellBender algorithm (remove-background)(v.0.2)[58] was applied to remove ambient and background RNA from each count matrix produced using the CellRanger pipeline. Downstream analysis was performed using the Scanpy package. Doublets with a score of *>* 0.15 were removed using Scrublet[59]. We performed quality control and filtering based on the following settings: number of genes *>* 500, total counts *>* 1000, % mitochondrial < 5 (nuclei), % mitochondrial < 20 (cell), % ribosomal < 5 (nuclei), % ribosomal < 20 (cell), red blood cell score < 1 (nuclei), red blood cell score < 2 (cell).

### 4.5 Single-cell data integration

After pre-processing and removal of low-quality cells and nuclei, highly variable genes were identified with `sc.pp.highly variable genes, flavor=’seurat’)`, retaining features with mean expression between 0.0125 and 3 and normalized dispersion ≥ 0.5, computed per batch (batch key=’batch key’), followed by integration using scVI[10, 11], accounting for categorical covariates of donor, cell or nuclei, and kit 10x, as well as continuous covariates of total counts, %mito, and %ribo. We then fit scVI using `scvi.model.SCVI(adata, n hidden=128, n layers=3, n latent=50, dispersion=’gene-batch’)` and trained for 50 epochs (`vae.train(max epochs=50)`), which yielded the embedding used for downstream analyses. Batch correction and bio-conservation were calculated using the scib-metrics python package[60]. Neighbourhood identification, high-resolution leiden clustering, and further dimensional reduction using UMAP were performed based on the scVI latent space. Additionally, Force Atlas or Diffusion Map embeddings were utilised for visualisation[61]. Subsequently, clustering was performed employing the Leiden algorithm. Cells and nuclei of extra-cardiac origin were removed based on marker gene expressions: lung epithelial cells (*IGFBP2, EPCAM, SOX2*, and *NKX2-1*); hepatocytes (*ALB, APOA1, TTR*); hepatic stellate cells (*CRHBP, FCN3, OIT3*); parathyroid cells (*PTH, MAFB, GATA3*); and further erythrocytes (*FTH1, HBA1, HBA2, HBB*). The remaining data were integrated using scVI with the same variables, followed by the downstream processing. To annotate cell types, cell barcodes comprising major clusters were separated from the initial raw count matrix for subclustering analysis and processed separately through the same downstream analysis as described. Subclusters were annotated based on immunohistochemical and spatial transcriptomics validation, the anatomical location sequenced after targeted dissection, and literature evidence.

### 4.6 Cardiac cell type abundance over time

Analysis was restricted to nuclei-derived libraries in the first trimester. To minimize sampling bias from targeted dissections, we excluded donors known to have targeted sampling (BRC2251, BRC2253, BRC2256, BRC2260, BRC2262, BRC2263, C104, C82, C85, Hst39, Hst40, Hst41). To focus on the heart proper and avoid variability introduced by inconsistent inclusion of great-vessel tissue across donors (i.e., different lengths or segments of aorta/ductus/pulmonary arches and pericardium being kept in or out), we removed cells known to pertain to these regions by excluding fine-grain annotations representing great-vessel and pericardial compartments (DuctusArteriosusSmoothMuscleCells, GreatVesselAdventitialFibroblasts, GreatVesselArterialEndothelialCells, GreatVesselSmoothMuscleCells, GreatVesselVenousEndothelialCells. To avoid sampling bias due to immune enrichment, and because immune nuclei could not be unambiguously attributed to cardiac versus great-vessel origin, we removed all immune-lineage nuclei (coarse-grain Leukocytes) and any FACS CD45^+^/CD45pos nuclei. For each week, we pooled nuclei across donors, counted nuclei per mid-grain lineage, and converted counts to proportions (lineage count divided by total nuclei that week). To present continuous trends while preserving local structure, weekly proportions were smoothed with a 3-point moving average (weights 0.25, 0.5, 0.25) and then interpolated with a shape-preserving PCHIP across the observed week range on a dense grid (n=300); negative interpolates were clipped to zero and proportions were renormalized at each grid point to sum to one. Stacked area charts and line plots were utilised for visualisation, through Matplotlib. Leukocyte abundance was plotted separately in the same manner, restricting the analysis to non-CD45 enriched samples.

### 4.7 Cell type correlation analysis

We quantified transcriptomic similarity among annotated cell types by plotting a cell-type correlation matrix computed in the modality/batch-corrected scVI latent space. Specifically, on the integrated AnnData object, we computed a dendrogram over fine grain categories with `sc.tl.dendrogram(adata, groupby=’fine grain’, use rep=’X scVI’)`, thereby ordering cell types by hierarchical clustering in the scVI embedding. We plotted a symmetric matrix of pairwise Pearson correlations between fine grain centroids in the scVI latent space (X scVI).

### 4.8 Gene set scoring

Gene set scoring was carried out using the `sc.tl.score genes()` function in Scanpy, with the default settings. Cell cycle genes[62], ribosomal genes, mitochondrial genes, genes related to hemoglobin, Y-chromosome genes were scored. *XIST* counts and the presence of Y chromosome gene expression was examined to determine the sex of the samples. Tip and stalk identity of capillary ECs were scored through a previously reported set of genes[63]. Cells with tip scores surpassing their stalk scores were identified as tip cells, while those with predominant stalk scores were annotated as stalk cells.

### 4.9 cell2location mapping

Cell2location[64] was used for mapping the suspension dataset on spatial transcriptomics slides; for a detailed description, see Cranley et al[9].

### 4.10 Differential expression and gene set enrichment analysis

Differential gene expression analysis was performed utilising the `sc.tl.rank genes groups()` function within the Scanpy framework. Differentially expressed genes were identified applying a significance threshold of p-value < 0.01 and a log2 fold change (log2FC) *>* 2. These identified gene lists were then subjected to exploration using Gene Ontology Biological Process 2023[65] through the GSEApy package[66].

### 4.11 Adult left right chamber comparison

We integrated our fetal dataset with adult (30 to 70 years old, donation-after-circulatory-death (DCD) cases) snRNA-seq from published human-heart single-cell atlases (PubMed IDs 35926050, 37438528 and 32403949). This was carried using using scVI with the same parameters as our fetal atlas as above. Exclusion of cells and donors was carried out, omitting cells of low-quality, stressed or contaminated cells, e.g., cells aberrantly expressing epicardial or fibroblast markers, notably concentrated in one study. Donors were excluded where any of the four chambers were missing and barcodes were retained when sourced from four cardiac regions, LV, RV, LA, RA. Left–right identity for CMs was assigned using our curated fetal annotations in fetal data and by anatomical sampling region (LV vs RV; LA vs RA) in adults.

Life periods were treated as an ordered categorical variable (’first trimester’ *>* ’second trimester’, *>* ’adult’), and analyses focused on two mid grain groups of the atria and the ventricles. For each life period and atria/ventricle, differential expression (DE) analysis was performed among the fine grain pertaining to that life period, using `sc.tl.rank genes groups()` with the Wilcoxon rank-sum test (`method=’wilcoxon’, key added=’wilcoxon’`). For each chamber group (L or R) within that period, we extracted DE results with `sc.get.rank genes groups df()` applying log2fc min = 1 (≥2-fold change) and pval cutoff = 0.01; filtering follows Scanpy’s implementation, which includes adjusted p-values (pvals adj). We then calculated, within the group, the fraction of cells expressing each candidate gene as the proportion of non-zero values in the working matrix, retained genes expressed in ≥10% of group cells, and used the resulting list both to summarize DE statistics. For scoring, within each life period we computed per-cell aggregate signatures for the L and R gene sets using `sc.tl.score genes()`, saved as correspondingLeftScore and correspondingRightScore. Each score vector was min–max normalised within the interval of [0,1], and written back to adata.obs. Lastly, a difference metric `Score Difference = correspondingRightScore - correspondingLeftScore` was calculated.

Pseudobulk analysis of left vs right cells at each stage was carried out using this dataset, and the adult heart cell atlas data Kanemaru et al., 2023[18]. Pseudobulk samples were generated by a single sample of a maximum of 100 cells from each chamber for each donor if available. Statistical tests were carried out using DESeq2[67] between conditions of Left vs Right with shrinkage of log-2-fold-changes using apeglm[68].

### 4.12 Coronary endothelium continuum

scFates[69] pipeline was employed to fit a curved trajectory across the embedded representation of the blood vessel endothelial cells. Genes that showed significant changes in expression over pseudotime were detected and their expression profiles were plotted. Among the coronary endothelial cells, the milestone linked to proliferating coronary endothelial cells was marked, and the associated features were plotted across the pseudotime trajectory.

### 4.13 Cell cell communication and niche analysis

We used CellPhoneDB to infer cell-cell interactions between annotated clusters[70]. Expression matrices and the metadata relating cell-barcode to cluster were submitted to CellPhoneDB. To compute niche interactions, we utilised the statistical method provided by the cpdb statistical analysis method function, 1000 iterations and applied a p-value threshold of 0.01. We further filtered the interactions by selecting the interactions where both the ligand and receptor expressions were present in more than 40 % of the cells within a particular cluster. Results were plotted using ktplots-py package[71].

### 4.14 Immunohistochemistry

Primary human fetal heart tissue was fixed overnight in 4 % PFA (Alfa Aesar) with gentle rocking at 4 °C. The next day, the tissue was incubated in 30 % sucrose (Sigma/Merck) in PBS overnight. On the following day, hearts were embedded in OCT (Sakura Tek), frozen on dry ice, then sectioned onto slides in 10 *µ*M thickness and kept in -80 °C until immunostaining. See Supplementary Table 9 for antibody details. For the staining of cTnT, SHOX2, and TBX3(**Fig. 2g**); PLA2G5, MYH11, PECAM1, and FOXC1 (**Fig. 4a**); TSHZ2, ADGRG6, and COL4 (**Supp. Fig. S8c, k**): Cryosectioned heart slides were initially thawed for 10 minutes at room temperature and then rehydrated in a tris-buffered saline solution. The sections were surrounded with a hydrophobic pen (Abcam) and subjected to permeabilization for 10 minutes in a buffer containing 0.25 % saponin (Alfa Aesar) in TBS. Following this, the permeabilization buffer was removed, and a blocking step was performed for 1 hour at room temperature using a blocking buffer, which was composed of 0.3M glycine (Merck), 10 % goat serum (Merck), and 0.2 % Tween-20 (Merck) in TBS. Subsequently, the blocking buffer was discarded, and the primary solution (Supplementary Table 9, in 10 % goat serum and 0.2 % Tween in TBS) was applied overnight at 4 °C within a humidified chamber. The samples underwent three 5-minute washes with a washing buffer containing 0.2 % Tween-20 in TBS. Secondary antibody staining (**Supplementary Table 9**) was carried out for 1 hour at room temperature. Following this, the slides were washed three times in the washing buffer before the application of DAPI-containing mounting medium (Vectashield). Finally, the sections were imaged using a Zeiss LSM 980 AiryScan microscope and analysed utilising Zeiss ZEN software.

For the staining of CNN1, ACTN2, and Phalloidin (**Fig. 2f**, **Supp. Fig. 6c**): Slides were hydrated in PBS for 15 min at RT. Slides were then permeabilised with 0.3 % Triton X-100 (Sigma Aldrich; X100)) in PBS for 15 min at RT. After washing with PBS, samples were blocked with 3 % BSA in PBS for 30 min at RT. Primary antibodies were then added in 3 % BSA in PBS and incubated overnight at 4 °C. After washing 3 times with PBS, the appropriate secondary antibodies and 1:1000 dilution of DAPI (Miltenyi Biotec; 130-111-570) for nuclear stain was also added in 3 % BSA in PBS for 2 hours at 4 °C. Cryosections were then washed 3 times with PBS before being mounted under a glass cover slip using Vectorshield Antifade Mounting Medium (Vector laboratories; H-1000-10). Imaging was then carried out using the Zeiss LSM 980 Airyscan2 confocal microscope. Images were processed using Fiji.

### 4.15 Quantification of CNN1 staining

Images from three sections were manually transformed. The left and right chambers were identified by the relative chamber wall thickness and the images were rotated accordingly. The compact myocardium in the bottom half of ventricles was traced and used as a mask for the analysis. Care was taken during the trace to avoid trabeculated tissue and the edge of the samples to avoid noise from coronary vasculature and other edge-artefacts. The green channel in this mask was extracted and overlayed against a red background, and the images imported into R using the package EBImage. Pixels were binned in squares, covering 16 bins across the horizontal axis to approximate 8 columns of RV, and 8 columns of LV. Mean intensity for each square bin was then calculated, omitting any pixels in the red channel. The average RV CNN1 intensity was calculated as the mean of all pixel bins in the first 8 columns while the average LV CNN1 intensity was calculated as the mean of all pixel bins in the latter 8 columns. To compare average LV vs average RV enrichment of CNN1 across the three donors, a paired Student’s t-test was used with an alpha of 0.05.

### 4.16 Embryonic stem cell-derived cardiomyocytes differentiation

Embryonic stem cells (RUES-2) are maintained in mTeSR Plus Medium (STEMCELL Technologies, USA). Before differentiation, cells are dissociated into single cells using TrypLE (Thermo Fisher, USA). Then, 900 × 10^3^ cells are seeded into each well of a 12-well plate using mTeSR Plus supplemented with 10 µM ROCK inhibitor Y-27632 (AOBIOUS, USA). The medium is then refreshed with mTeSR containing Matrigel (1:60; Corning, USA). On day 1 of differentiation, cells are cultured in RPMI 1640 (Gibco, USA) supplemented with 50X B-27 minus insulin (Thermo Fisher, USA) and 6 *µ*M Chiron-99021 (Universal Biologicals, UK). On day 2, an equal volume of RPMI 1640 with B-27 minus insulin is added. On days 3-4, cells are cultured in RPMI 1640 with B-27 minus insulin and 2.5 *µ*M Wnt-c59 (Universal Biologicals, UK). From day 5-6, cells are maintained in RPMI 1640 with B-27 minus insulin. From day 7-12, the cells are cultured in RPMI 1640 with B-27 containing insulin (Thermo Fisher, USA). Typically, beating cardiomyocytes are observed by day 10. On day 13, cardiomyocytes are dissociated into single cells using the STEMdiff Cardiomyocyte Dissociation Kit (STEMCELL Technologies, USA). Then, 1 × 10^6^ cells are seeded into each well of a 12-well plate in RPMI 1640 with B-27 plus insulin supplemented with 10 *µ*M Y-27632. On days 14-17, cardiomyocyte purification is performed using RPMI 1640 without glucose (Thermo Fisher, USA), supplemented with 4 nM sodium L-lactate (Sigma-Aldrich, USA) and 1:100 MEM Non-Essential Amino Acids (Gibco, USA). From day 18 onwards, cardiomyocytes are maintained in RPMI 1640 with B-27 containing insulin.

### 4.17 PRRX1 knockdown

The effect of PRRX1 siRNA (Thermo Fisher, USA) and Silencer Select Negative Control No. 1 siRNA (Thermo Fisher, USA) is evaluated using hPSC-CMs from day 20–25. Cells are incubated with 30/45/60/75 nM siRNA and DharmaFECT siRNA transfection reagent (1:500; Horizon Discovery, UK) in RPMI 1640 with B-27 plus insulin for 48 hours. The hPSC-CMs are then incubated with refreshed 48 nM siRNA for another 48 hours. The medium is then replaced with fresh RPMI 1640 with B-27 plus insulin for an additional 24 hours before cell lysates are collected. RNA extraction is performed using the GenElute Mammalian Total RNA Miniprep Kit (Sigma-Aldrich, USA). RNA quantity and quality are assessed using NanoDrop (Thermo Fisher, USA). cDNA synthesis is conducted with the PrimeScript RT Reagent Kit (Takara Bio, USA). Quantitative polymerase chain reaction (qPCR) is performed using QuantStudio 5 Real-Time PCR (Thermo Fisher, USA) and SYBR Green (Thermo Fisher, USA), with 3 biological replicates from independent differentiations and transfections. Primers for PRRX1 (TGATGCTTTTGTGCGAGAAGA and AGGGAAGCGTTTTTATTGGCT) and 6 right vCMs transcriptional markers including *ANKRD1* (AGTAGAGGAACTGGTCACTGG and TGTTTCTCGCTTTTC-CACTGTT), *CNN1* (CTGTCAGCCGAGGTTAAGAAC and GAGGCCGTC-CATGAAGTTGTT), *CYP2J2* (TCCATCCTCGGACTCTCCTAC and GTCAC-CAAGCTCCAAGCTAAAA), *GPNMB* (AAGATTGCCACTTGATGCCG and TCC-CTCATGTAAGCAGAAGGTC), *ZNF385D* (CTTCCCCAATTTCAATGCGATG and CAACTGGCAAATGTTGCATGATA), and *KCNIP4* (AATGCCCCAGTGGT-GTTGTTA and TGAAATCCTCGAAACTCACAGC) are used. Relative gene expression was calculated using the Delta-Delta Ct Method with GAPDH as endogenous control. Comparison between *PRRX1* siRNA and scramble siRNA groups were performed using paired T-test. p-value <0.05 was considered statistically significant.

### 4.18 RNAScope analysis

Simultaneous detection of *HES5*, *HEYL*, *DLL4* mRNA with MYH11 protein was performed on frozen sections using Advanced Cell Diagnostics (ACD) RNAscope®2.5 LS Multiomic core reagent Kit (Cat No. 322930), RNAscope Multiomic C1, C2, C3, and C4 Channel reagents (Cat No. 322935, 322940, 322945, and 322950 respectively) plus RNAscope®2.5 LS Probes (Supplementary Table 9) (ACD, Hayward, CA, USA), in combination with ACD Protease free reagent from the RNAscope kit (ACD, Hayward, CA, USA), and MYH11 antibody (Sigma Aldrich, Cat: HPA015310). Briefly, sections were removed from the -70 °C freezer, immediately placed into pre-chilled 10 % NBF and fixed for 1 hour at 4 °C before rinsing in 70 % ethanol and then 100 % ethanol x2 for 3 mins each. Slides were air dried before loading onto a Bond RX instrument (Leica Biosystems). Heat pre-treatment with Epitope Retrieval Solution 2 (Cat No. AR9640, Leica Biosystems) at 95 °C for 8 minutes, before a protease-free reagent step for 30 minutes at ambient. Probe hybridisation, signal amplification and detection were performed on the Bond Rx according to the ACD protocol. Probes were visualized using Opal fluorophores (Opal 570, Opal 620, Opal 690 and Opal 780; Akoya Biosciences Cat No. FP1488001KT, FP1495001KT, FP1497001KT, and FP1501001KT, respectively) diluted using RNAscope LS Multiplex TSA Buffer (see table). MYH11 protein detection was performed with primary antibody diluted 1:500 in Bond Antibody Diluent (Leica, Cat No.AR9352) for 15 minutes at ambient, followed by incubation with the Polymer-HRP component of Leica’s Polymer Refine Detection System (Leica, Cat No. DS9800) and Opal 520 (Akoya Biosciences Cat No. FP1487001KT) detection at 1:1000 dilution. Slides were counterstained with DAPI from the RNAscope kit according to manufacturer’s instructions. Slides were then removed from the Bond Rx and mounted using Prolong Diamond (ThermoFisher Cat No P36965). The slides were imaged on the Zeiss AxioScan Z1 to create whole slide images. Images were captured at 40x magnification, with a resolution of 0.11 microns per pixel and analysed with QPath software.

### 4.19 Curation of datasets for CMageR

Several datasets were used for training or external validation of both CellTypist model and mGAM, including 4 published datasets[1, 8, 16, 39] and 3 sampled datasets from our atlas of covariate-balanced variations of cells or nuclei (Supplementary Materials). For Farah2024[16], the data were curated at fine grain for reporting accuracy by atlas-level integration with using scVI our dataset and another endothelial dataset[72], followed by manual annotation to harmonise labels. To ensure impartiality here, no label transfer methods were used and the workflow relied only on prior metadata and a cross-referencing of gene markers. The held-out data contained no cells annotated as Pericardial cells, Leukocytes, or a subset of mural cells, presumably resulting from different sampling and dissociation strategies. To align our predictions back with the author’s original annotations, our fine grain labels were mapped hierarchically back into the appropriate cell type. For Cui2019[1] and DeBono2025[39], cells were annotated manually using author annotations to fit within our labelling scheme. For RvCM vs LvCM labels, we gathered the dissected region information from published metadata and curated an anatomical ground-truth[1, 16] for left and right vCM identity. Where dissected region information was absent, we utilised given annotations from published data[39]. The cell type labels for external datasets using in both training and held-out test for the Cell Age models were annotated by mapping the author’s original annotations into our coarse grain or mid grain level annotations depending on the model granularity.

### 4.20 CellTypist model training

The CellTypist models were trained on our dataset only by assembling a carefully balanced sample of cells and nuclei (10x scRNA-seq and 10x multiome snRNA-seq) data from all 21 hearts within this study. This resulted in a training dataset of 22,813 cells or nuclei (**Supp. Fig. S10a**). The summary of training dataset variables as well as barcodes and metadata are found in **Supplementary Table 5**. For the final CMageR cell type models, three independent CellTypist models were trained on our sample using the standard CellTypist model training pipeline[29]. Specifically we used the `celltypist.train()` function in python on cells and nuclei normalised by library size (counts per 10,000 reads). Features for the CellTypist model were not pre-screened and one model was generated for each granularity without additional optimisation. For assessing generalisability and cross-modality performance, additional independent CellTypist models were constructed. In these models, performance was evaluated on held-out donors from our dataset as well as on an external held-out dataset Farah2024[16], which comprises whole-cell transcriptomes only. Prior to training, datasets were restricted to the intersection of shared genes, library-size normalised (10,000 counts per cell), and log-transformed, ensuring a consistent feature space across all splits and modalities.

### 4.21 Cardiac clock training

The `CMageR` developmental age mGAMs were trained on a combined sample of snRNAseq barcodes from hearts in our dataset (11 hearts), and barcodes from scRNA-seq hearts from an additional two external datasets[8, 16], totalling 33 donor hearts of mixed cells and nuclei. The number of cells used to train each model was different depending on the cell type while care was taken in sampling to balance the contributions from each donor within each cell type and thus a partial balancing of developmental time (**Supplementary Table 5**). For the snRNA-seq within our data, the fine grain annotations were also balanced across time prior to joining the datasets to avoid spurious temporal latent effects of cell type heterogeneity or technical biases across the time points (see example mid grain CMs sample in **Supp. Fig. S14a**). One time-course model was trained for each cell type in our dataset where data were sufficient. Cell types with insufficient data were mid grain PericardialCells, or fine grain PericardialCellsFibrous, PericardialCellsParietal, Monocytes, MonocytesMPOpos, DendriticCellsMature, MastCells, NaturalKillerCells, and ProBCells.

#### 4.21.1 Feature Selection

Feature selection for each cell type was carried out through several steps (Supplementary Materials). Genes were initially screened for dynamic behaviour across developmental time in our snRNA-seq dataset only (*hearts = 11*) by fitting a univariate GAM on gene expression as response, and PCW as the predictor using mgcv[73] (`gam(gene` ∼ `s(week, k = 6, bw = "cr"), data = data, method = REML)`), or as a linear model (`lm(gene` ∼ `week, data = data)`). Smoother functions were estimated by residual maximum likelihood (REML). Significant gene trajectories were then tested for linearity by comparing the smoother fit to a linear model using Akaike Information Criterion (AIC). Gene smoothers were predicted and derivatives were calculated (51 time points between 4 and 20 PCW) to determine where periods of significant change occur using methods from Simpson et al., (2018)[74] (Supplementary Materials). Confidence intervals were then defined for each point and significant slopes were determined with an alpha of 0.05. Non-linear genes were then further categorised manually into early, mid, late, overall, or long time bins depending on when their derivative was significant, and sorted by absolute log-fold change (See Supplementary Materials), resulting in a total of 17 categories of genes with varied significant developmental time trajectories (**Supp. Fig. S12a**, **Supplementary Materials**). The top genes were taken from the screening categories and were filtered further to remove those within a curated list of technical noise features such those differentially expressed between cells or nuclei or biological sex (Supplementary Materials). As a final step in feature generation, the binary effect of cells or nuclei was included as an additional fixed effect term `modality`, where interactions were also explicitly included for every gene to accommodate for observed gene-specific mixed modality effects between cells and nuclei.

#### 4.21.2 model training

An mGAM was trained for each cell type using PCW as the response variable and gene expressions of the pre-screened genes, modality, and the interaction terms as predictors using the function `cv.gamsel(x, y, family="gaussian", gamma=0.5, …)`[31]. The feature maximum degrees were pre-specified to the rounded-up effective degrees of freedom (EDF) parameter in each gene’s `mgcv()` fit to a maximum of 4, while each gene × modality interaction EDF was constrained to EDF - 1. The dfs parameter was set equal to the degree but with a maximum of 3 to constrain overfitting. The degree and dfs parameters for the fixed-effect of modality were both set to 1. The models were trained using 11 non-overlapping folds, each with 3 independent hearts in a L3HO cross-validation where the regularisation parameter λ and final model fits were selected using the 1-standard-error rule[31]. This forms the final model for each cell type of mid grain and coarse grain annotations. For the fine grain cell types, `cv.gamsel()` was run using a different strategy on 11 hearts from our dataset only (See Supplementary Materials). Out of fold performance was then assessed for a conservative assessment of performance in the coarse grain, and mid grain models by re-creating 11 individual models for each cell type using the same feature space and architecture as the final trained model. However, the cross-validation used 10-folds (30 hearts) while a single fold (3 hearts) was completely omitted from the training and cross-validation step.

#### 4.21.3 alternative models

Alternative models were constructed using the usingthe single-cell architecture surgery approach outlined in scArches[30]. Three models were built as an additional independent mapping method for confidence in the LvCM and RvCM findings (Supplementary Materials). One model was constructed to compare alternative models for mapping cardiac developmental time. In each case, an scVI[11] model was created using our all of, or subsets of our dataset, and a label transfer approach scANVI[32] was used to build a discrete classification model for the classes of interest (fine grain cell types, left vs right vCMs or other CMs, CM mid grain time bins). Query datasets (in vitro, or in vivo) were then integrated and annotated using the trained models.

### 4.22 Temporal gene over-representation analysis

Overrepresentation analysis (ORA) was carried out using Gprofiler2[75] on all genes in the VentricularCardiomyocyte mGAM. At each time point (n = 51), all genes with a significant positive slope, and all genes with a significant negative slope were collected (*p < 0.05*). Each time point’s significant genes were queried against a background of genes expressed in the vCM cluster regardless of age (at least 3 counts in at least 3 vCM barcodes). This created 51 sets of results for increasing GO:BP terms, and 51 sets of results for decreasing GO:BP terms, one set of terms for increasing genes and one set of terms for decreasing genes at each smoother time point. Then, in each of increasing, or decreasing sets, the terms were filtered by adjusted p value (*p < 0.01*) and ranked by each GO:BP’s maximum F-score across the time period. This was calculated using precision as the fraction of query genes within the term genes, and recall as the fraction of term genes within the query. By retaining non-significant representations for each process throughout the entire period for all terms, we were able to plot the p-value and F-score for all terms at all time points.

### 4.23 Stem cell model benchmarking

For all of the hPSC-CM datasets and the *in vivo* datasets, we acquired processed counts matrices from publicly available repositories as linked in the original articles[40–49]. These matrices contained varied feature sets due to processing and alignment differences between publications. Predictions were made independently in each dataset to avoid a minor intersection of features. Predictions were made using the steps outlined in the CMageR pipeline through CellType prediction, hierarchical annotation, and age-prediction. Reported cell types in the results were made using the filtered predictions for a more confident interpretation of the *in vitro* models.

### 4.24 Data availability

Raw sequencing data from the single-cell RNA-seq and single-nucleus Multiome RNA-seq datasets are available at the European Nucleotide Archive (ENA)(https://www.ebi.ac.uk/ena) at EMBL-EBI, under accession ID PRJEB115510. The raw Visium data from our sister paper[9] is located at ENA under accession ID PRJEB76689. All of the processed data are available for browsing gene expression and download from the Heart Cell Atlas (https://www.heartcellatlas.org/foetal.html, publicly available at the time of publication). We also developed a summary-level interactive data explorer on R shiny for this dataset accessible at https://www.sinha-lab.org/resources-1. The human reference genome (GRCh38) used for read mapping is available from 10x Genomics (https://support.10xgenomics.com/single-cell-gene-expression/software/release-notes/build).

### 4.25 Code availability

The code used for all the analysis performed in this study and sister study[9] is available on GitHub at https://github.com/sinhalab-csci/Developing_heart_atlas/ and in part at https://github.com/Teichlab/Developing_heart_atlas. The `CMageR` pipeline python package is available at GitHub (https://github.com/KniSche/cmager/).

## 5 Acknowledgements

We thank the donors for granting access to the tissue samples. We also thank Roger Barker for his help in coordinating samples. We thank staff at the Wellcome Sanger Cytometry Core Facility, Cellular Genetics Informatics team and Core DNA Pipelines team for their support; A.Oszlanczi for her help on sample management; B. Ç akır for his help on the Heart Cell Atlas web portal; and A.Wilk for administrative assistance. This work was made possible by a partnership between the Wellcome-MRC Cambridge Stem Cell Institute at University of Cambridge and Wellcome Sanger Institute. We acknowledge that some samples in this work were provided by the MRC/Wellcome Trust-funded Human Developmental Biology Resource (HDBR). We thank Heather Etchevers and team at INSERM for their courteous communication regarding our findings on their *in vivo* dataset samples.

This project was made possible in part by the BBSRC strategic Longer Larger grant CellTalkHHD (S.A.T., S.S., R.T., V.K.S., K.K., and M.K.K); the British Heart Foundation (BHF) Senior Fellowship (FS/18/46/33663) (S.S. and L.G.); the Chan Zuckerberg Foundation (2021-237882 to S.A.T.); Wellcome Trust (WT206194 to S.A.T); S.A.T. is also funded by the CIFAR Macmillan Multi-scale Human Program; the Oxbridge BHF Centre for Regenerative Medicine (RM/17/2/33380) (V.K.S.); the Overseas Research Fellowship of the Takeda Science Foundation to K.K; Wellcome Trust Clinical PhD Fellowship to J.C.; and BHF grants PG/17/24/32886 (L.G.) and RG/17/5/32936 (H.D.). We also acknowledge core support from the Wellcome Trust, the Medical Research Council and the Wellcome Trust–Medical Research Council Cambridge Stem Cell Institute. This research was funded, in whole or in part, by the Wellcome Trust (grant no. 203151/Z/16/Z).

## 6 Author contributions

V.K.S. - Project design, Analysis (Annotation, Modelling, LR identity, CMageR, summary web app), Manuscript; S.B. - Project design, Analysis (Annotation, communication, spatial validation, LR identity, adult comparison), Manuscript; J.C. - Project design, Data generation, Analysis (Annotation), Manuscript; K.K. - Project design, Data generation, Analysis (Annotation), Manuscript; B.W. - immunohistochemistry; C-K.W. - PRRX1 knockdown; H.D. - heart collection, processing, and sequencing; M.C., J.C.M.L., M.K.K., and R.T. - Critical input into design and conception; M.D’S. - additional CellTypist model training; K.P. - Analysis (Visium mapping); L.R., C.I.S., R.K., S.P. - assisted snRNA-seq data generation; M.D., I.M., M.P. - data generation (Visium); S.Y.H. - assisted with anatomical structure annotation; X.H. - data generation (tissue procurement); L.G. - heart collection, processing and sequencing, provided critical input into the design; S.A.T. and S.S. - overall design and supervision.

## 8 Supplementary Figures

**Supplementary Figure S1.**
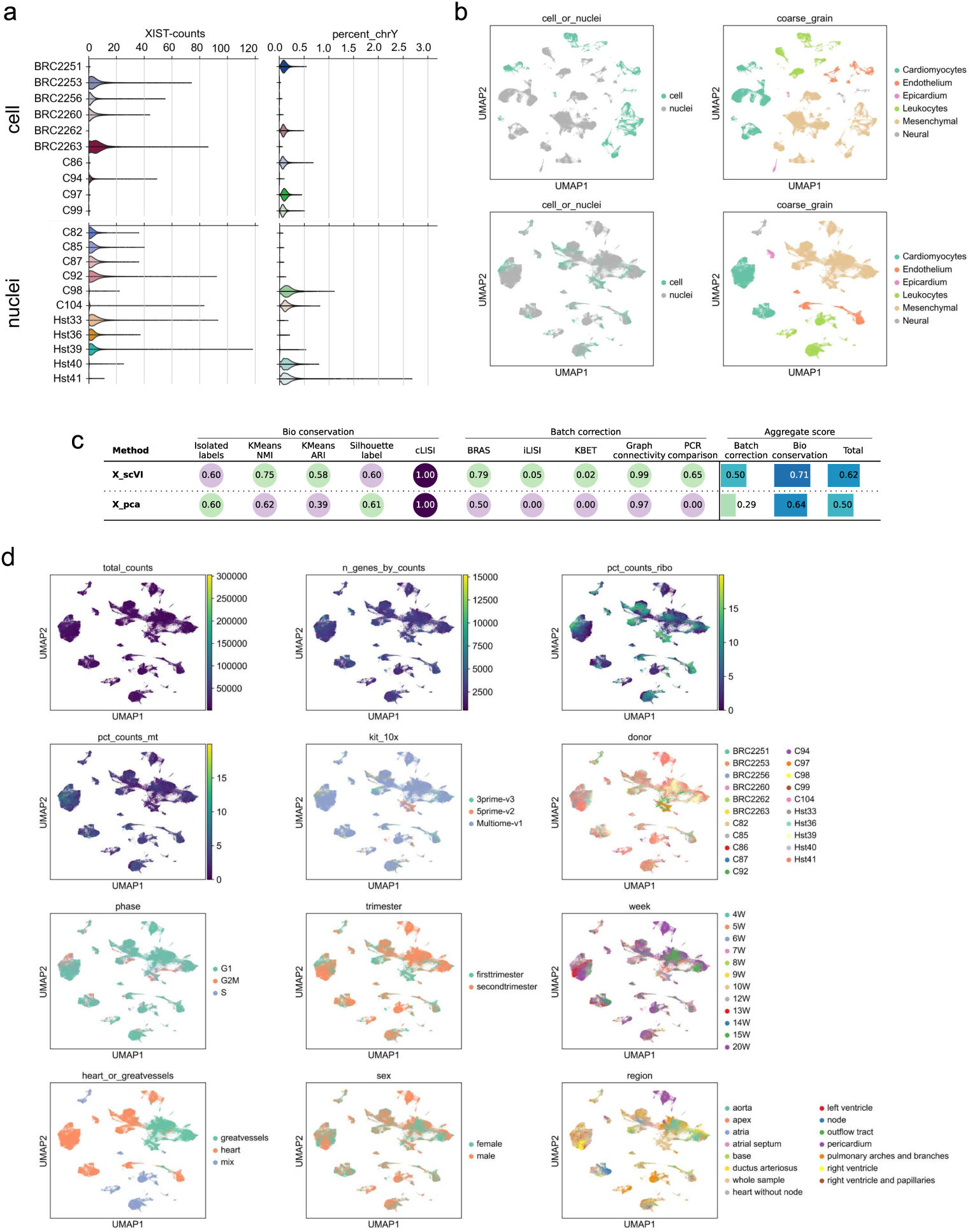
**A:** Sex inference of the samples. XIST counts and Y-chromosome counts in single-cell (upper panels) and single-nuclei (lower panels) processed samples. **B:** Integration of cell and nuclei data using scVI. UMAPs display cell vs nuclei labels (left) and coarse-grain annotations (right). scVI integration (lower panels) aligns cells and nuclei of the same type into unified clusters, preventing modality driven separation seen without integration (upper panels). **C:** Batch correction for “batchkey”, and bio-conservation metrics for “coarse grain” before (x pca) and after (x scVI) batch correction using scVI. Calculated using scib-metrics. **D:** UMAPs of the global embedding colored by technical and biological covariates. Shown are total counts, number of detected genes, mitochondrial and ribosomal percentages, 10x kit, donor, cell cycle phase, developmental stage (trimester, week), location after annotation (heart vs great vessel), sex, and dissection region.

**Supplementary Figure S2.**
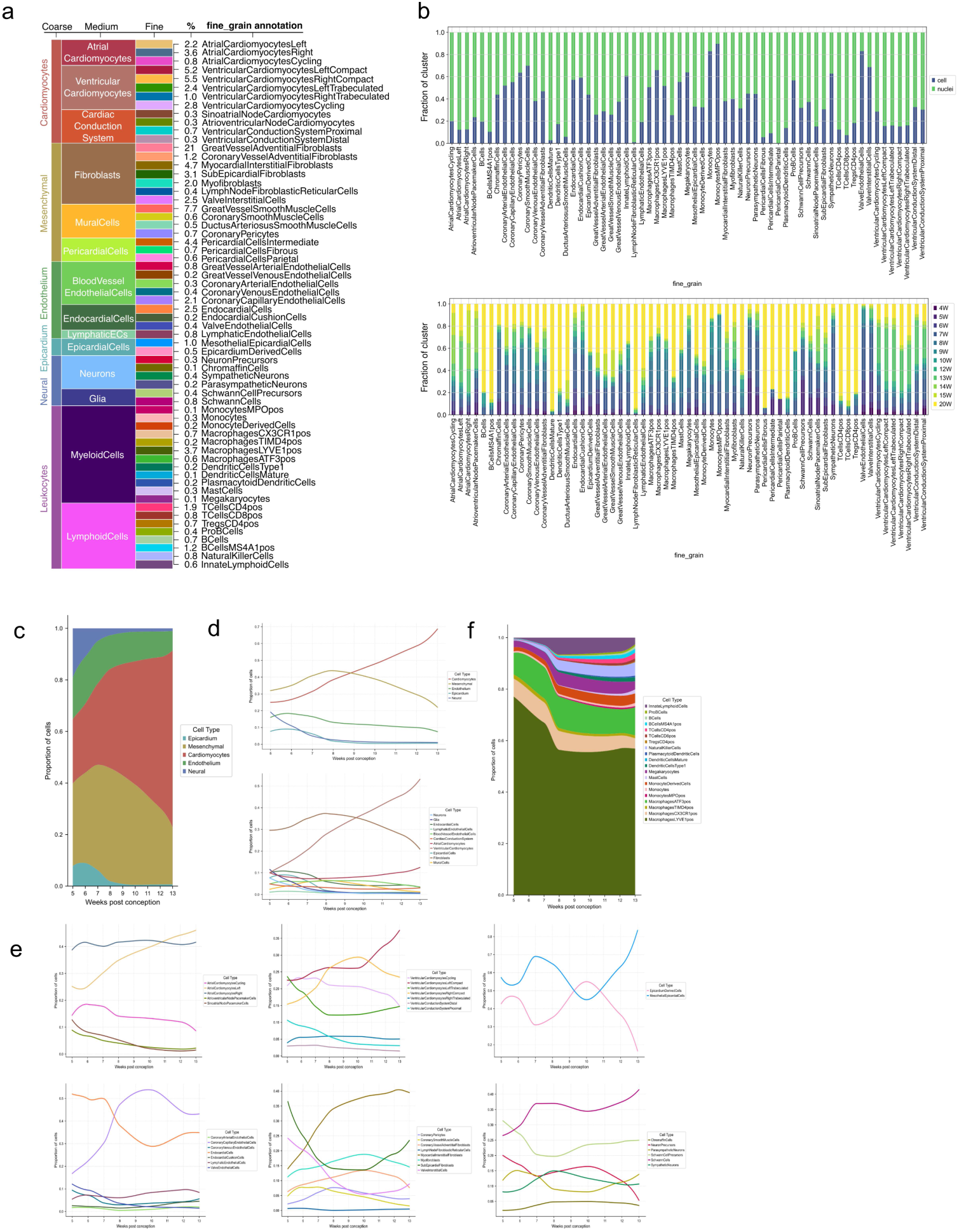
**A:** Hierarchical cell type annotation. Shown are coarse, medium and fine-grain categories, with percentages in the dataset provided for each fine-grain. **B:** Distribution of fine-grain cell types based on modality (cell vs nuclei, top) and developmental stage (weeks, bottom). **C, D, E:** Changes in cell type composition of the human heart during the first trimester, shown as proportions. To reduce sampling bias, immune cells (CD45 enriched), great vessel cells and pericardial cells were excluded due to varied dissection across samples. Only whole hearts rather than targeted dissections were included: c Coarse grain cell type dynamics shown as stacked proportions; d, line plots of coarse- and mid-grain categories; and e, line plots of fine-grain subtypes grouped by their respective higher level mid-grain categorisation. **F:** Fine grain dynamics of immune cell populations shown separately as stacked proportions. This analysis only includes whole heart-processed samples.

**Supplementary Figure S3.**
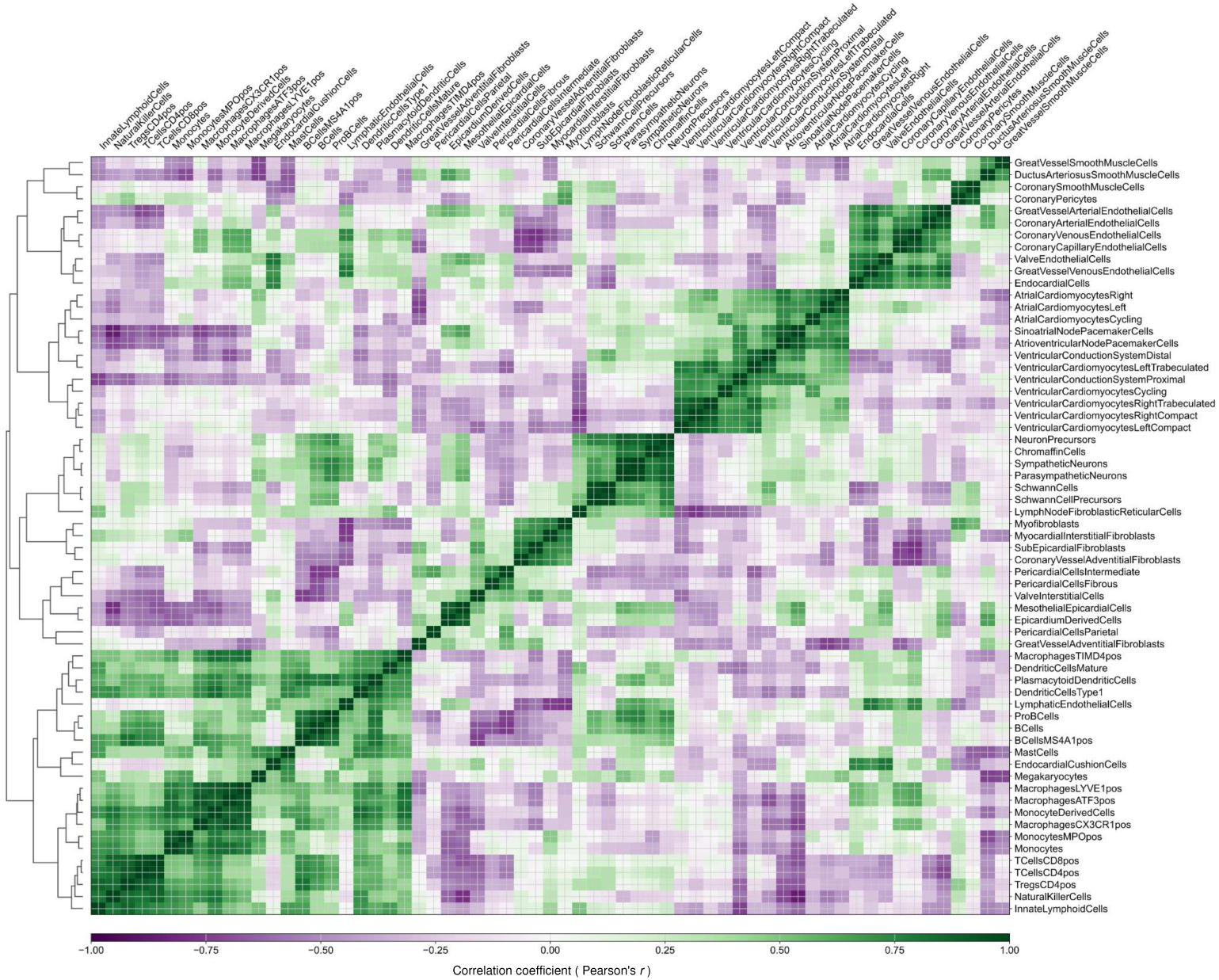
Matrix plot of pairwise transcriptomic similarity among fine grain cell types. Pairwise Pearson correlations were computed between celltype centroids in the modality/batch-corrected scVI latent space and rows/columns were ordered by hierarchical clustering (from the precomputed dendrogram). The heatmap is scaled from -1 to 1, with green indicating positive and purple indicating negative correlation. Block patterns along the diagonal highlight clusters of transcriptionally related populations.

**Supplementary Figure S4.**
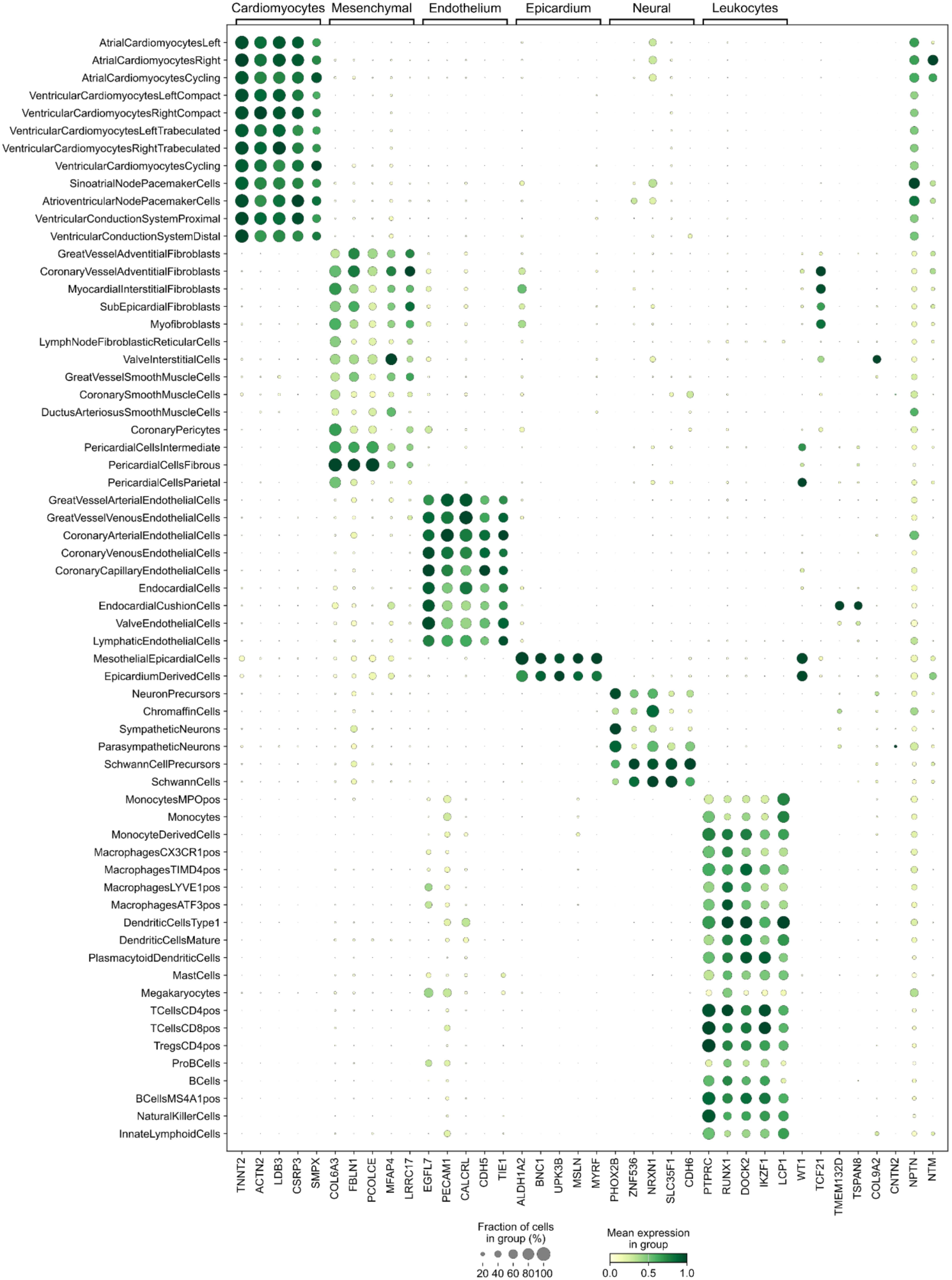
Marker gene expression across the atlas. Dot plot showing differentially expressed markers associated with each coarse-grain cell label. Each dot represents the expression of a marker gene in a given fine-grain population. Dot size indicates the fraction of cells within the population expressing the gene, while dot color represents the mean expression level, scaled from 0 (lowest expression of that gene across all cell types) to 1 (highest expression of that gene across all cell types). The plot also includes several genes referenced throughout the article on the right. The same dot size and color scaling scheme is applied to all dot plots in this paper.

**Supplementary Figure S5:**
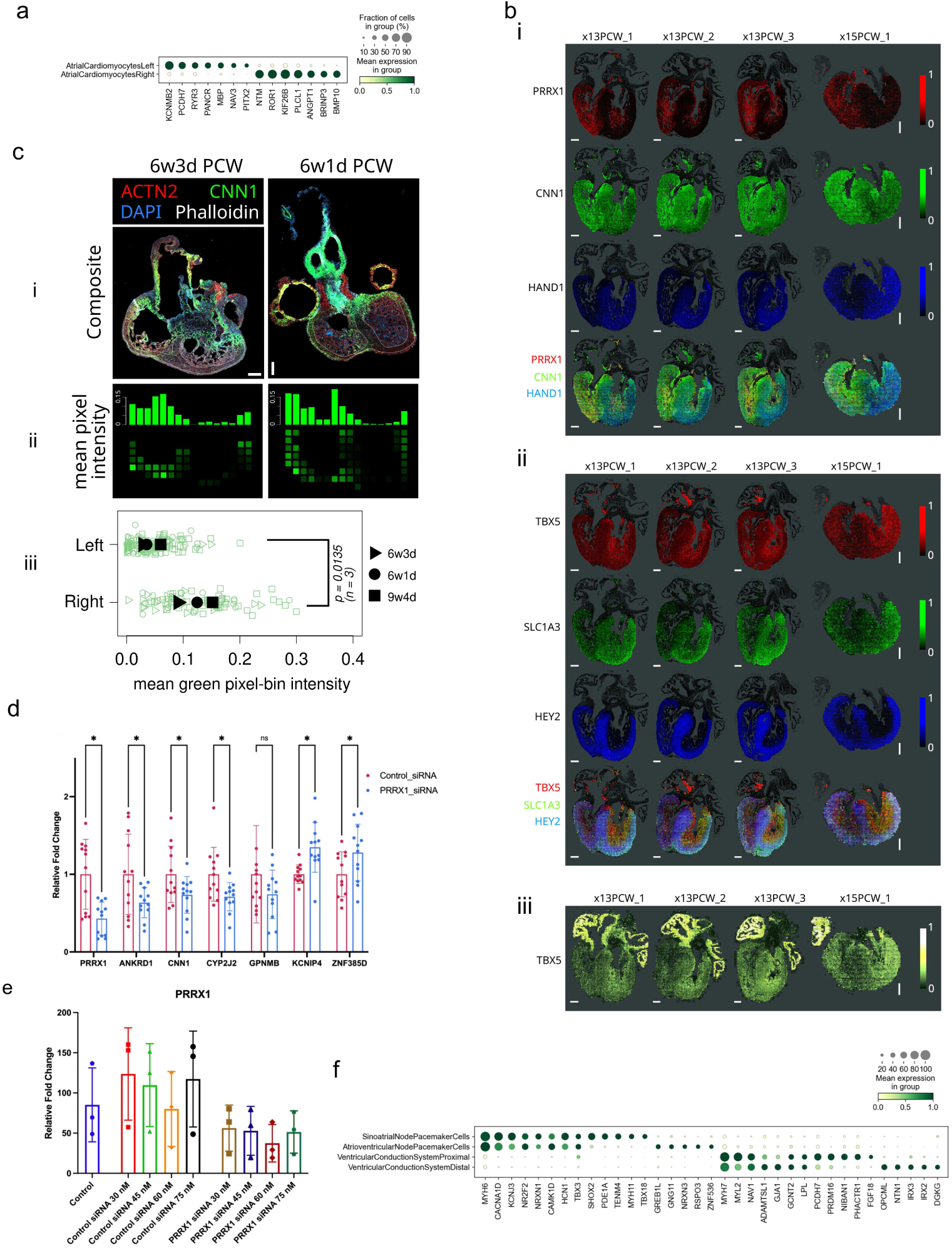
Left-right transcriptomic differences in atrial and ventricular cardiomyocytes and conduction system markers. **A:** Dot plot depicting the top markers differentially expressed between the right and left atrial cardiomyocytes. **B:** Multiplexed Error-Robust Fluorescence in situ Hybridization (MERFISH) spatial transcriptomics data from 4 human fetal hearts shows i, and ii, left and right separation across selected left and right ventricular cardiomyocyte markers; and iii, *TBX5* the putative first heart field marker. Values show mean expressions calculated from cells in 100 *µ*m square bins using data accessed from Farah et al., 2024[16]. Scale bar is 1 mm. **C:** Extended protein-level validation of higher right ventricular *CNN1* in human fetal hearts shown as i, immunofluorescent staining for CNN1 expression (scale bar: 200 *µ*m); ii, quantification of mean binned pixel intensities in outlined compact ventricular layer; and iii, comparison of mean binned pixel intensities across all three biological replicates stained for CNN1 comparing bins in the first 8 columns (right ventricle) against those in the last 8 columns (left ventricle) (*Student’s t-test, n = 3*). **D:** PRRX1 knockdown in embryonic stem cell-derived cardiomyocytes. PRRX1 knockdown was associated with downregulation of the right compact ventricular cardiomyocytes transcriptional markers *ANKRD1*, *CNN1* and *CYP2J2*. Conversely, *KCNIP4* and *ZNF385D* were upregulated. Data represented results from different biological replicates with independent differentiation and transfection. Asterisk (*) indicate statistical significance *p-value < 0.05*. **E:** Trial of different siRNA concentrations and the effect on PRRX1 knockdown. Consistent but *PRRX1* knockdown was achieved using siRNA at 30, 45, 60, and 75 nM concentration. Each data point represents individual biological replicate. Asterisk (*) indicate statistical significance *p-value < 0.05*. **F:** Dot plot depicting the top markers differentially expressed between the cells of the cardiac conduction system.

**Supplementary Figure S6.**
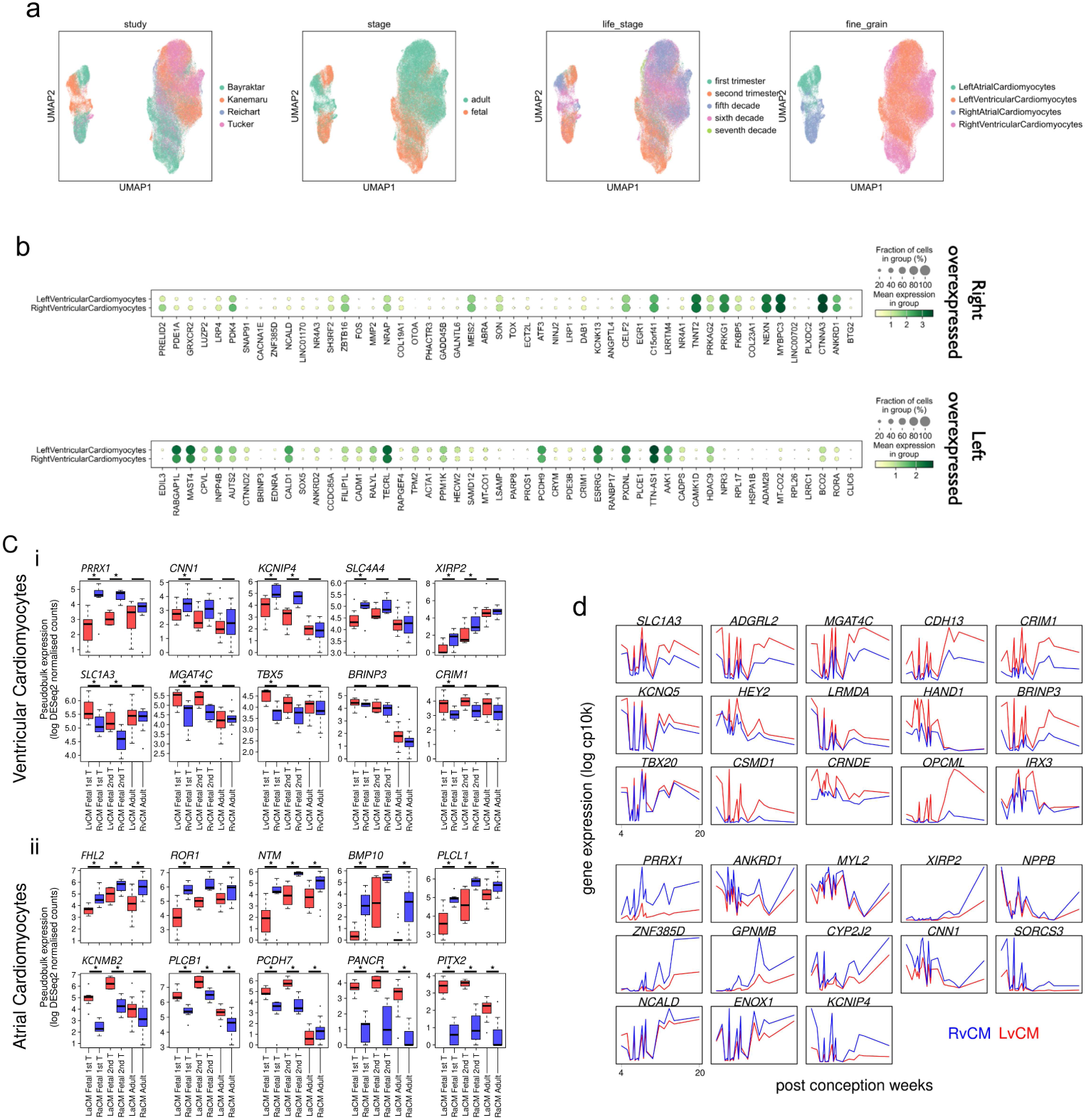
Left-right transcriptomic differences in atrial and ventricular cardiomyocytes continued. **A:** Uniform manifold approximation and projection (UMAP) embeddings of atrial cardiomyocytes (aCMs) and ventricular cardiomyocytes (vCMs) in single-cell RNA-sequencing integrated from fetal (this study) and adult reference datasets (see methods). **B:** Dot plot of top differentially expressed genes in adult vCMs. Genes most enriched in RvCMs (top) and LvCMs(bottom) defined by anatomical dissection are shown. Despite being identified as top DE genes in their respective groups, most display very similar expression profiles in both right and left cardiomyocytes, highlighting the modest transcriptional differences between right and left adult vCMs. **C:** Expression patterns of selected left and right differentially expressed developmental genes in i, vCMs and ii, aCMs across donor hearts as pseudobulk samples in the first trimester, second trimester, and adult our dataset (Fetal), and in adult data (Kanemaru et al 2023)[18]. Asterisk (*) denotes significance when (adjusted p <0.05, DESeq2). Note *CNN1* is **D:** Left and right vCM marker gene expressions between either LvCM (red) or RvCM (blue) clusters *within each donor* are consistent across the developmental window. Expression values depict the log counts per 10,000 UMIs (cp10k) as means of the the respective RvCM or LvCM clusters.

**Supplementary Figure S7.**
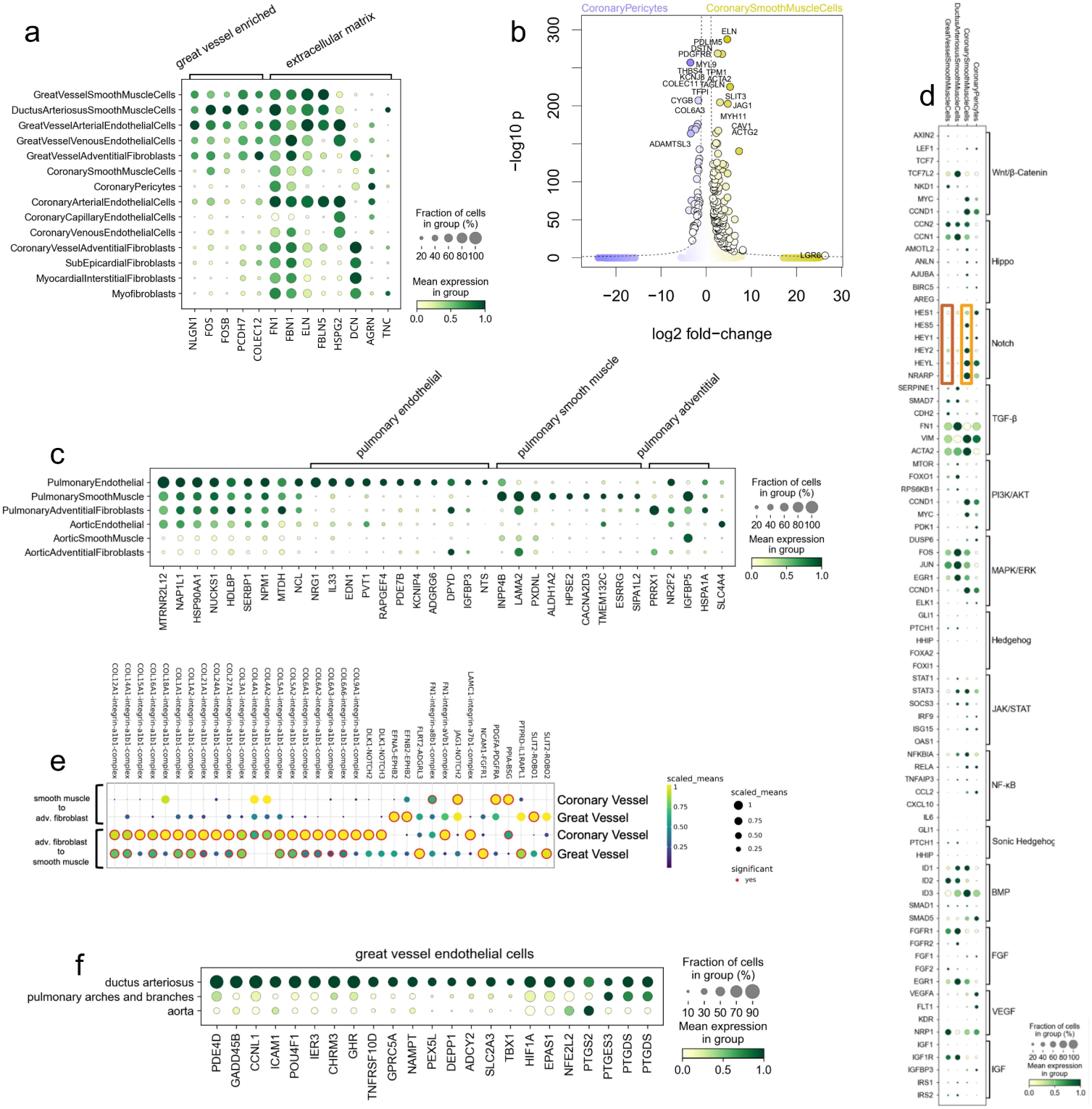
Comparative characterization of coronary and great vessel populations. **A:** Genes enriched in and shared across the great vessel constituents for great vessel and coronary vessel comparison, alongside the expression profile of various extracellular matrix proteins. **B:** (Volcano plot of the differentially expressed genes between coronary pericytes and coronary smooth muscle cells (CSMCs). **C:** Pulmonary artery specification of great vessel endothelial cells (ECs), SMCs and adventitial fibroblasts, through comparing the cells obtained from the targeted dissection of the aorta and pulmonary artery. **D:** Dot plot showing the expression of signalling pathway mediators across mural cell populations. Notch target genes display differential expression between coronary and great vessel SMCs. **E:** The interactions between SMCs and adventitial fibroblasts of the coronary and great vessels at the media-adventitia interface. **F:** Expression profile of the ECs obtained from the targeted dissection of ductus arteriosus, compared to the ECs obtained from the targeted dissection of the aorta and pulmonary artery of the same donor.

**Supplementary Figure S8.**
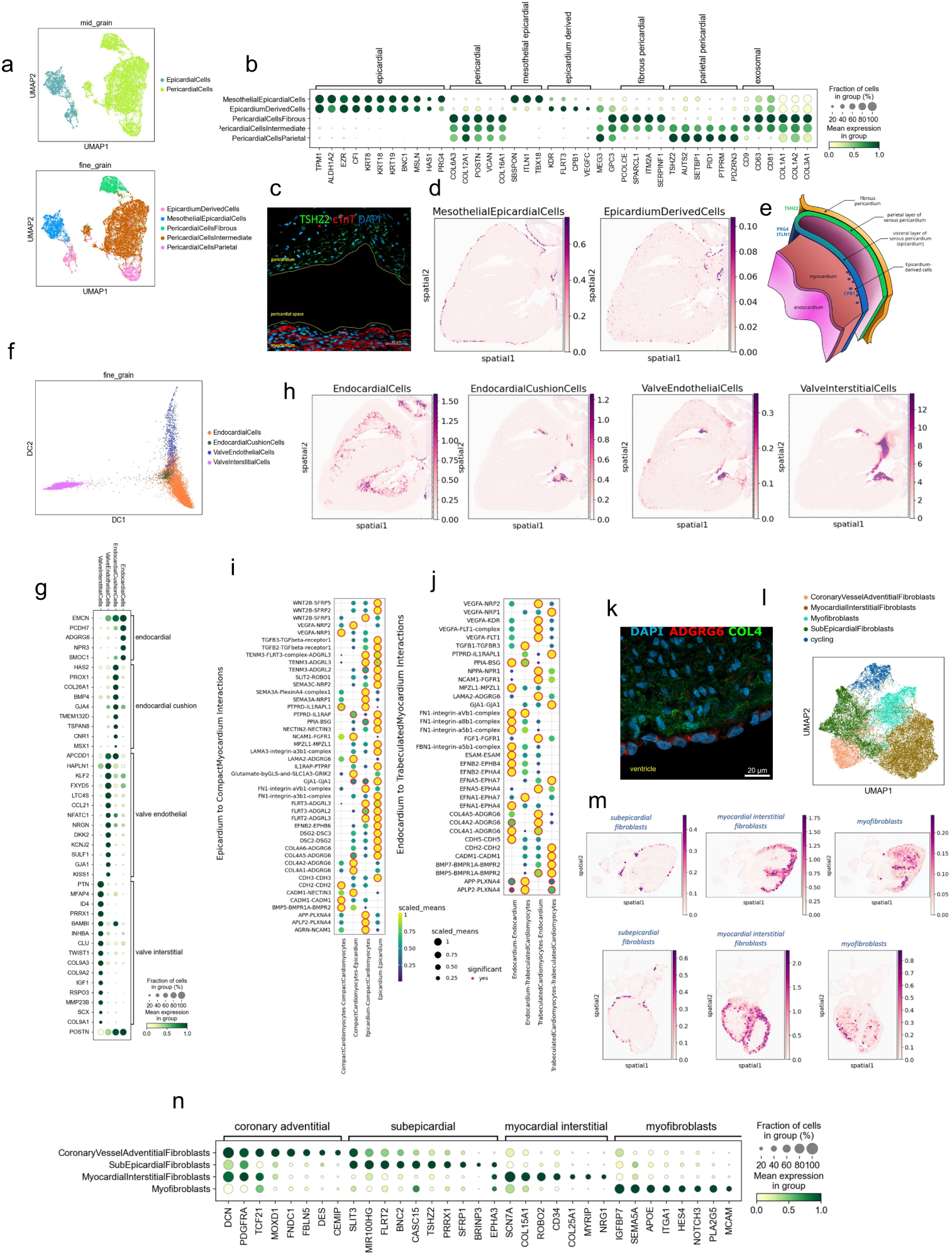
Pericardium, endocardium and myocardial fibroblasts. **A:** Two-dimensional uniform manifold approximation and projection (UMAP) embeddings of epicardial and pericardial fine-grain labels based on gene expression from cells in all samples and all stages. **B:** Dot plot displaying the differentially expressed genes associated with epicardial and pericardial cell types. **C:** Immunohistochemical analysis validates expression of the TSHZ2 at the parietal layer of the serous pericardium. **D:** cell2location spatial mapping of mesothelial epicardial cells and epicardium derived cells on a 16 PCW spatial transcriptomics section. **E:** Schematic illustrating the cell types comprising pericardium and the marker genes associated with them. **F:** Diffusion map embedding of the endocardial and valve cell types. **G:** Dot plot displaying the differentially expressed genes associated with endocardial and valve cell types. **H:** cell2location spatial mapping of endocardial and valve cells on a 16 PCW spatial transcriptomics section. **I:** The cell-cell interactions between the epicardium and the cardiomyocytes of the compact myocardium. **J:** The cell-cell interactions between the endocardium and the cardiomyocytes of the trabeculated myocardium. **K:** Immunohistochemistry validation of expression of ADGRG6 at endocardium and COL4 at myocardium. Scale bar, 20 M. **L, M:** Fibroblasts of the heart shown as l, clusters on UMAP; and m, with distinct transmural localisation patterns mapped by cell2location. **N:** Dot plot displaying the differentially expressed genes associated with heart fibroblasts.

**Supplementary Figure S9.**
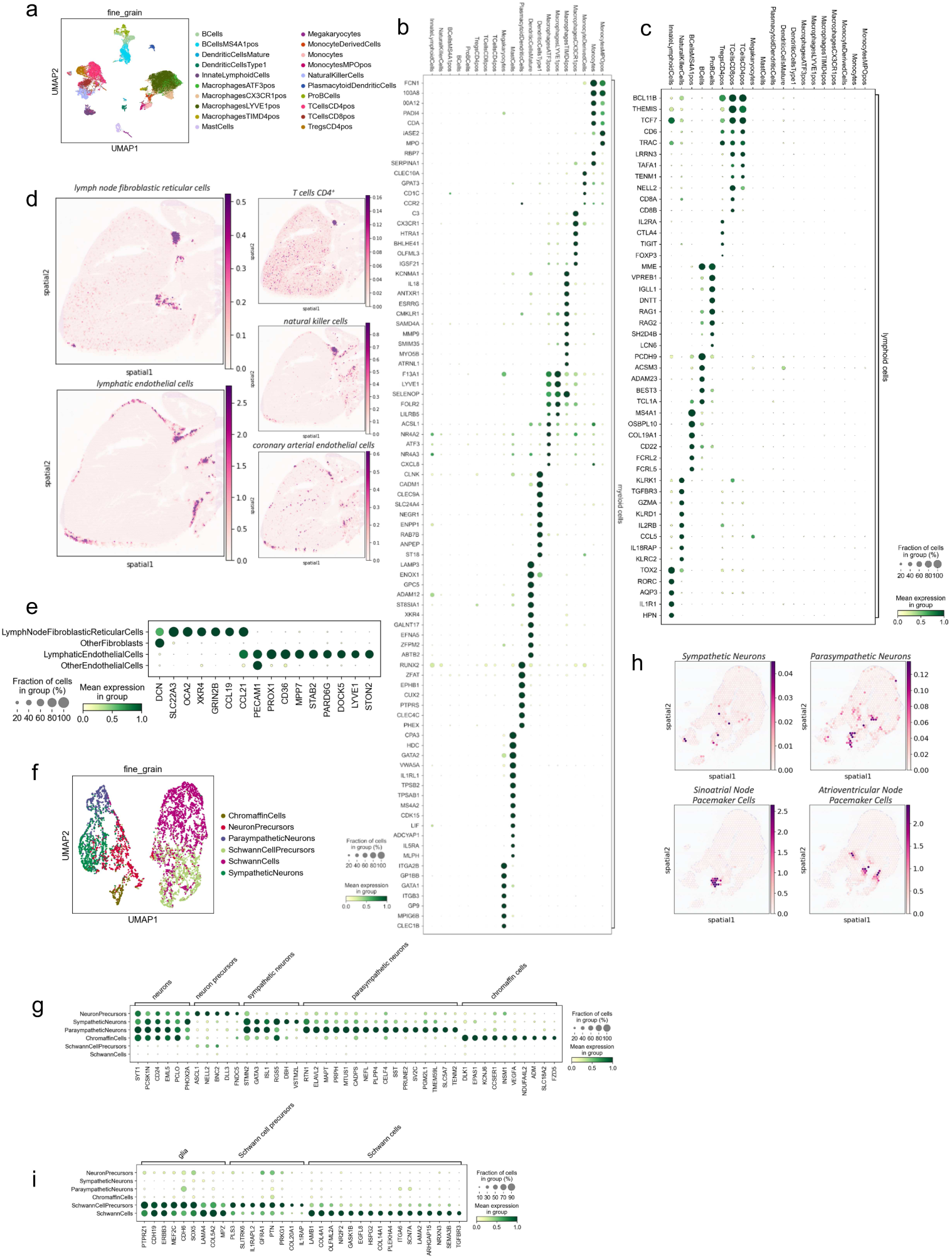
Leukocytes, lymphatic cells and neural cells. **A:** Subclustering of the coarse grain Leukocytes cluster reveals a diversity of fine grain annotated immune cells across the uniform manifold approximation and projection (UMAP). **B:** Differential gene expression analysis for the mid grain myeloid cluster with results shown as a dot plot for across all fine-grain cell types. **C:** Differential gene expression analysis for the mid grain lymphoid cluster with results shown as a dot plot for across all fine-grain cell types. **D:** cell2location mapping of lymph node fibroblastic reticular cells and lymphatic ECs on Visium slides localises these cells on a cardiac lymph node, alongside other leukocytes. **E:** Dot plot depicting the differentially expressed genes of lymph node fibroblastic reticular cells and lymphatic ECs. **F:** Cells of the neural compartment. **G:** Dot plot displaying the differentially expressed genes associated with neurons. **H:** cell2location mapping of sympathetic and parasympathetic neurons on Visium slides localises these cells in proximity to nodal pacemaker cardiomyocytes. **I:** Dot plot displaying the differentially expressed genes associated with glia.

**Supplementary Figure S10.**
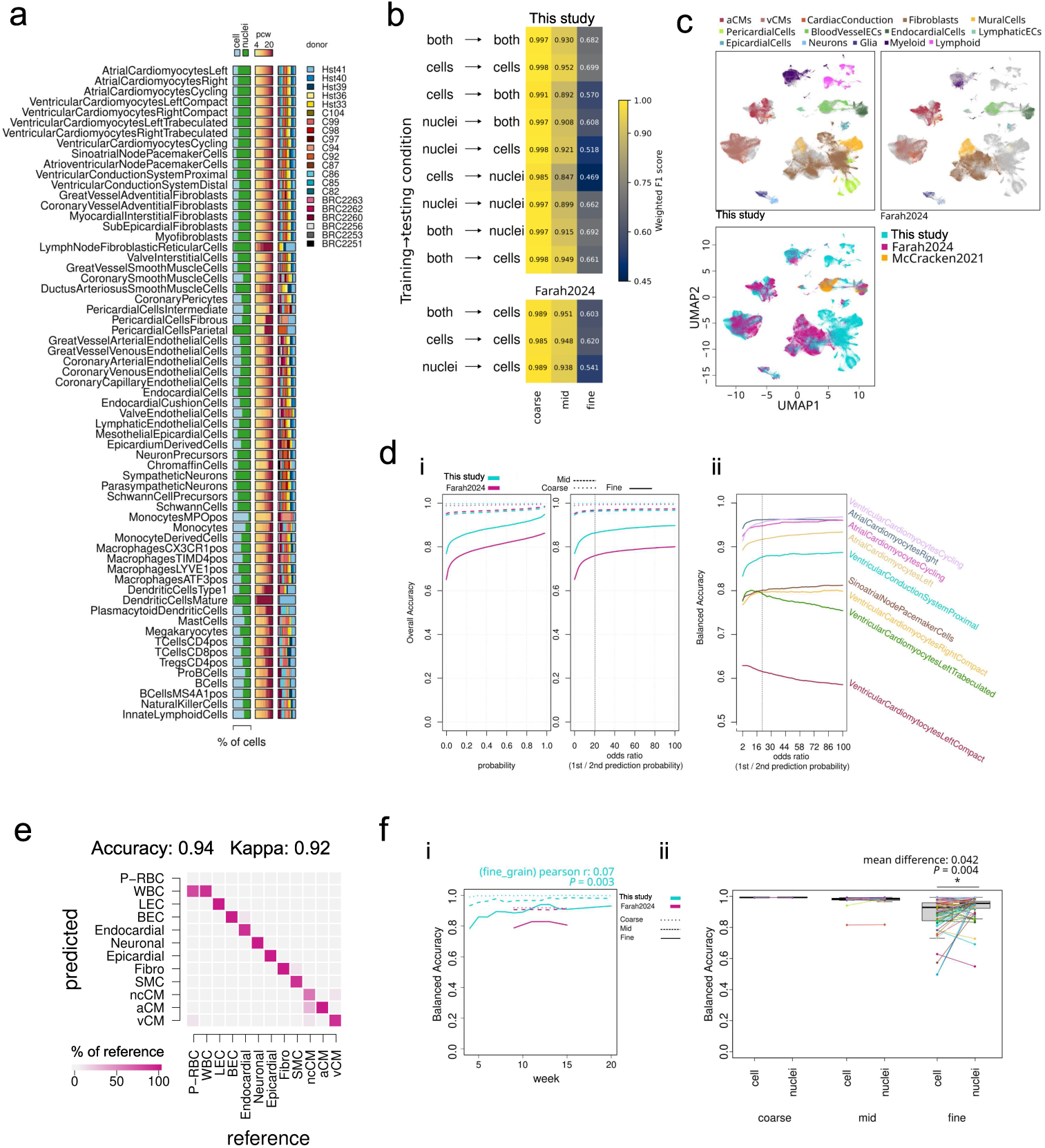
CellTypist Model sampling, validation, and statistics. **A:** Sample populations of technical variables across discrete classification CellTypist model training dataset 1 showing cells and nuclei balance, age balance, and donor balance within cell types at the fine grain resolution. **B:** Reduced generalisability of the CellTypist model was seen when trained only on cells, or nuclei, in contrast with a model trained on both. Models were trained on non-overlapping folds of train, held-out test, and then external held-out data. **C:** Integration of primary held-out scRNA-seq dataset Farah2024 and additional dataset McCracken2001 shows the congruence in manually re-annotated mid grain labels between datasets through uniform manifold approximation and Projection (UMAP). **D:** i, curve showing effect on overall accuracy when applying an increasing probability threshold; or ii, an odds-ratio between first and second prediction probability threshold to filter out predictions in coarse, mid, and fine grain annotations across test (Bayraktar2025) and validation (Farah2024) datasets. **E:** Confusion matrix and accuracy when collapsing cell type fine grain predictions into the original author annotations in Farah2024. **F:** Balanced accuracy of the coarse grain, mid grain, and fine grain CellTypist models; i, over time in test (This dataset) and validation (Farah2024) data (statistics calculated with Pearson’s correlation over mean accuracy for each week, n = 12); and ii, within cell and nuclei subset test data (statistics calculated using Welch’s two-sample t test, n = 63).

**Supplementary Figure S11.**
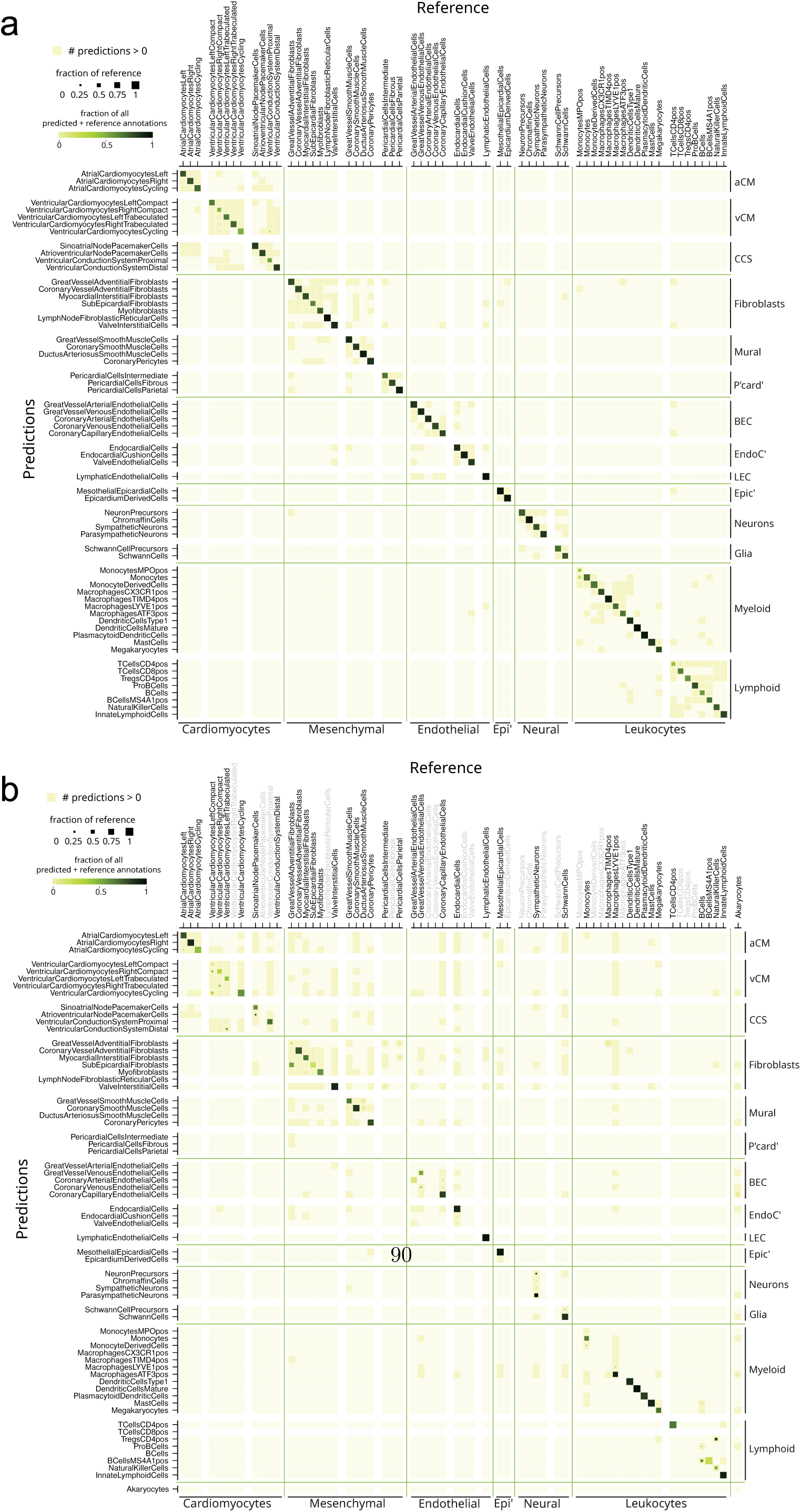
CellTypist fine grain confusion matrices. **A:** Confusion matrix of cell type classification on all fine grain predictions on validation barcodes - snRNA-seq or scRNA-seq barcodes that were not used to train the model but were from the same hearts. **B:** Confusion matrix for held-out scRNA-seq dataset Farah2024. These hearts were completely unseen by the model but annotations were manually curated by integrating the datasets, re-clustering, and using canonical gene markers. Annotations in lighter grey illustrate cell types not re-annotated explicitly in clusters in the validation dataset and their statistics against reference labels could not be calculated.

**Supplementary Figure S12.**
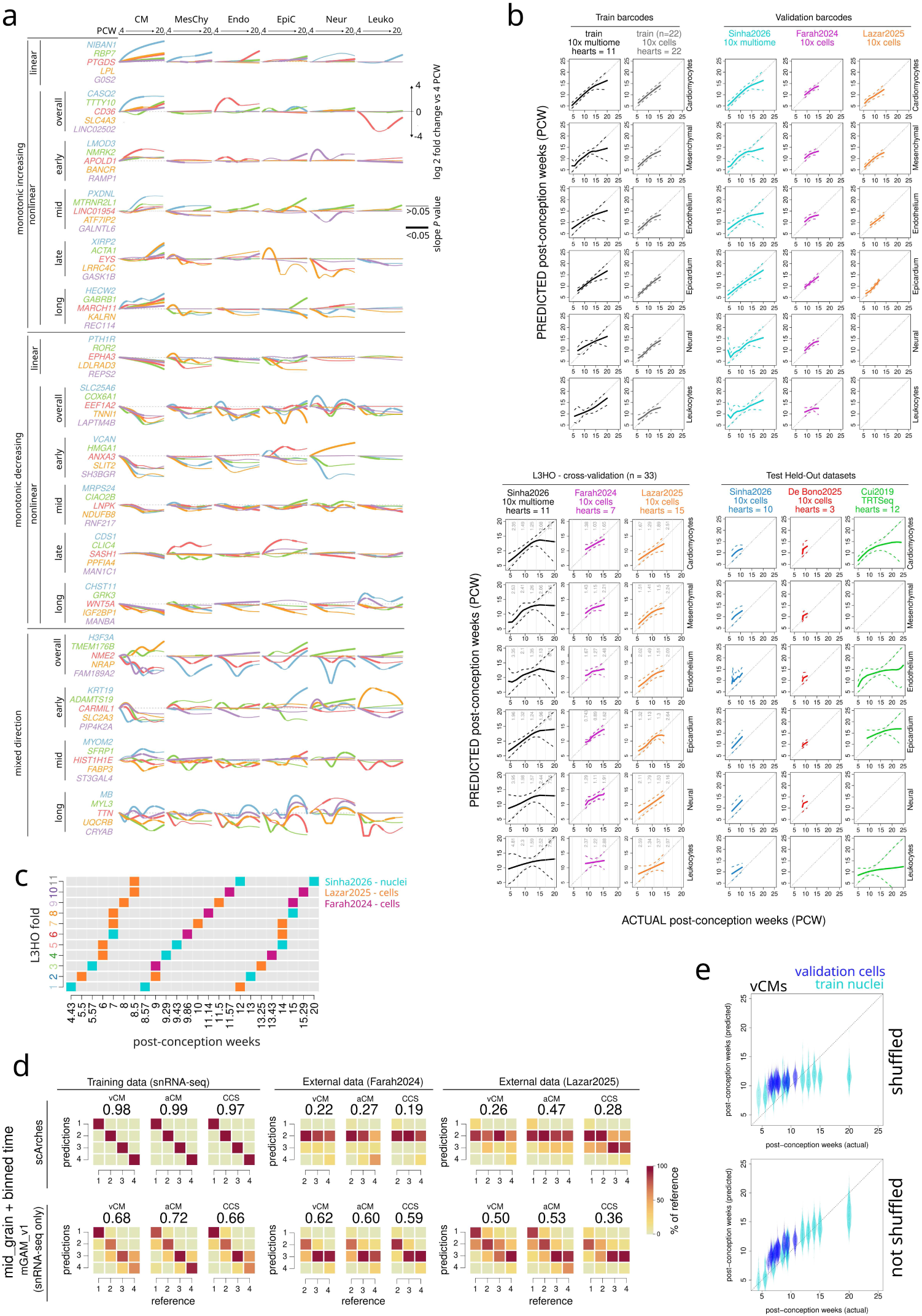
Extended assessment of cell age model. **A:** Modelling balloon plots of the top 5 Cardiomyocyte gene trajectories in each of 17 shape categories across all coarse grain cell types illustrate the cell type specificity needed for predicting cell age from transcriptional information. The tonicity and time-windows were predefined to balance features as input into the multivariate generalised additive model. Note, no “late” time-binned mixed-direction dynamic shapes were categorised in the Cardiomyocyte model. The y-axis shows the log 2 fold change of each gene from its expression at 4 post-conception weeks (PCW), with PCW as time on the x-axis. Columns separate the different coarse grain cell types. **B:** Final average age prediction fits of all coarse-grain cell types in training barcodes, validation barcodes, leave-3-heart-out (L3HO) out-of-fold cross-validation, and held-out hearts shown as smoothed mean predicted age for each heart (solid line) and the RMSE (dotted lines). Time-binned RMSEs are shown for L3HO cross-validation curves. **C:** The distribution of datasets and time points spread across the 11 L3HO folds. **D:** Comparison of current model methods with a standard atlas reference mapping method using single-cell architecture surgery with scArches and scANVI. Individual multivariate generalised additive models (mGAMs) were trained on our balanced sample of single nuclear RNA-seq barcodes (snRNA-seq) from our dataset (hearts = 11) only, one for each of ventricular cardiomyocytes (vCMs), atrial cardiomyocytes (aCMs), and cardiac conduction system (CCS) cells (see methods). A reference scVI model was trained on all snRNA-seq CMs (n = 11 hearts) from our dataset with a 12-class scANVI model for vCM, aCM, and CCS split into four different time windows (1: 4 to 7 PCW, 2: 7 to 10 PCW, 3: 10 to 14 PCW, and 4: 14 to 20 PCW). **E:** Predicted age vs Actual age in mid grain vCMs in validation snRNA-seq barcodes (turquoise) and held-out test scRNA-seq (blue) when shuffling or not shuffling the cell type models.

**Supplementary Figure S13.**
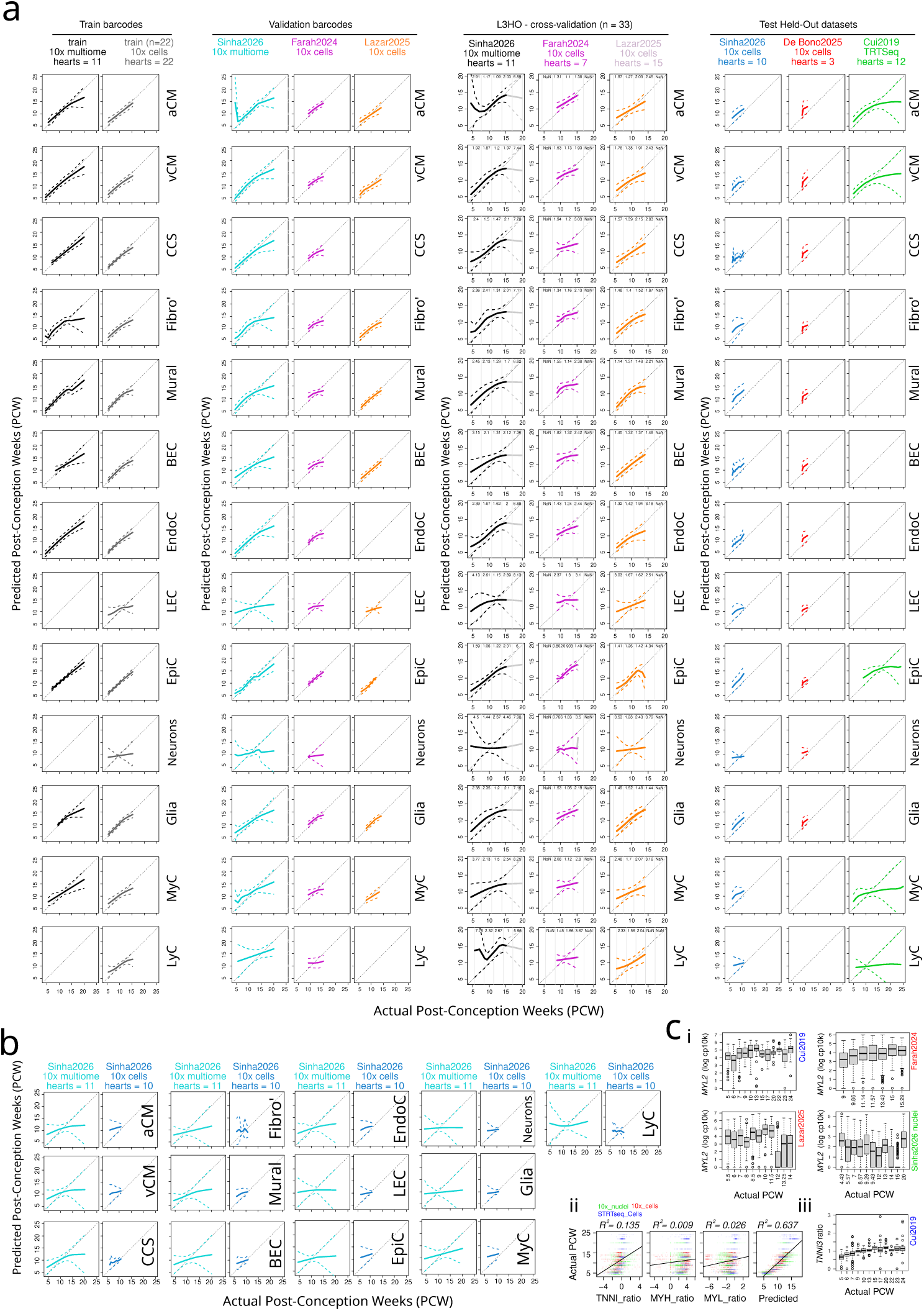
Cell age model fits in mid grain cell types. **A:** Final average age prediction fits of all mid grain cell types in training barcodes, validation barcodes, leave-3-heart-out (L3HO) out-of-fold cross-validation, and held-out hearts shown as smoothed mean predicted age for each heart (solid line) and the RMSE (dotted lines). Time-binned RMSEs are shown for L3HO cross-validation curves. **B:** Predicted age in mid grain cell types after randomising which cell type model is used on each cell. Datasets shown here are smoothed mean predictions (solid line) and RMSE (dotted lines) of validation single-nuclear RNA-seq barcodes (barcodes from the same hearts using in training, but not the same barcodes), and held-out test single-cell RNA-seq barcodes. **C:** Standard ratiometric measures of cardiomyocyte (CM) maturation are shown to be noisy and inconsistent predictors of cardiomyocyte age between 4 and 25 post-conception weeks (PCW): i, *MYL2* expressions in mixed datasets; ii, simple linear model fits and R-square value of actual PCW in response to predictors of log *TNNI3:TNNI1* ratios, log *MYL2:MYL7* ratios, log MYH7:MYH6 ratios, and the Predicted PCW from our model in ventricular CMs (vCMs) (*n = 2843*); and iii, the plateau in expression of *TNNI3* after 15 PCW.

**Supplementary Figure S14.**
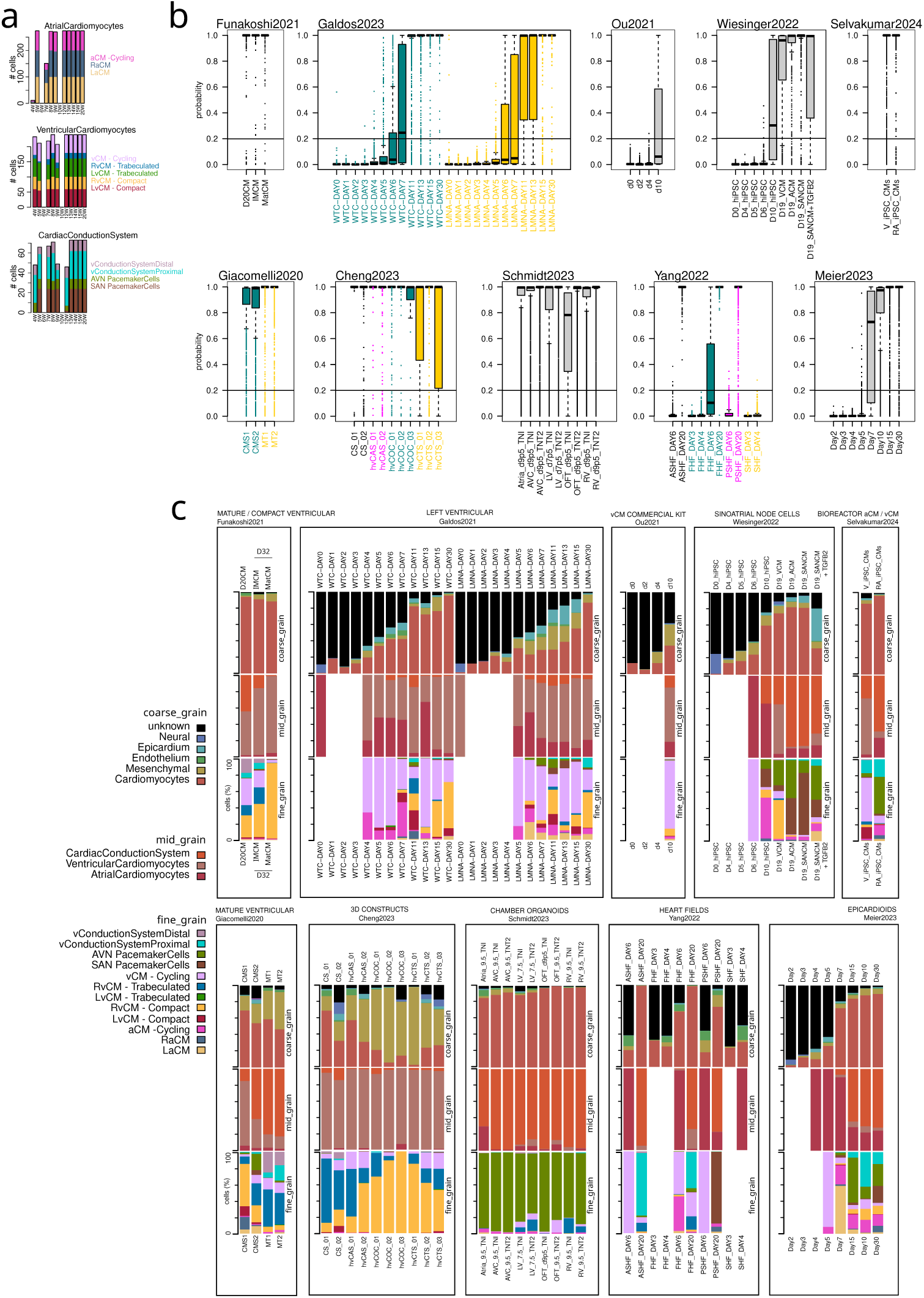
Extended hPSC-CM cell type results. **A:** Cardiomyocyte (CM) fine grain population balances within each time-point within the training dataset for each CM mid grain multivariate generalised additive model. **B:** CM prediction probability across all conditions in all datasets for CM labelled cells using CellTypist. The horizontal line shows the heuristic threshold of 0.2 for confidently calling CMs from the stem cell model data. **C:** Full suite of predictions for the 10 *in vitro* model datasets showing population fraction of different cell types, Including coarse grain predictions of all cells including fraction of “unknown” classified cells, CM-only filtered mid grain predictions, and CM-only filtered fine grain predictions. Study condition glossary: Selvakumar2024 - V iPSC CMs (Ventricular induced pluripotent stem cell cardiomyocytes), RA iPSC CMs (Retinoic Acid+ induced pluripotent stem cell cardiomyocytes); Galdos2023 - WTC (WTC cell line), LMNA (SCVI-111 cell line); Giacomelli2020 - CMS* (cardiomyocyte only replicate), MT* (microtissue replicate); Ou2021, d0, d2, d4, d10 (days 0, 2, 4, & 10); Funakoshi2021 - D20CM (day 20 cardiomyocytes), IMCM (day 32 - maturation media), MatCM (day 32 + maturation media); Cheng2023 - CS (cardiac spheroids replicate), hvCAS (human ventricular cardiac anisotropic sheet replicate), hvCOC (human ventricular cardiac organoid chamber replicate), hvCTS (human ventricular cardiac tissue strip replicate); Wiesinger2022 - d0, d4, d5, d6, d10, d19 (days 0, 4, 5, 6, 10 & 19), VCM (day 19 ventricular cardiomyocytes), ACM (day 19 atrial cardiomyocytes), SANCM (day 19 sinoatrialnode cardiomyocytes), SANCM+TGFB2 (day 19 sinoatrialnode-transitional zone cardiomyocytes); Schmidt2023 - d7p5 & d9p5 (days 7.5 & 9.5), AVC (atrioventricular canal), LV (left ventricle), RV (right ventricle), OFT (outflow tract); Yang2022 - FHF(first heart field), SHF (secondary heart field), ASHF (anterior secondary heart field), PSHF (posterior secondary heart field) Legend glossary: aCM (atrial cardiomyocytes); vCM (ventricular cardiomyocytes); AVN (atrioventricular node); SAN (sinoatrial node); R* (right); L* (left).

**Supplementary Figure S15.**
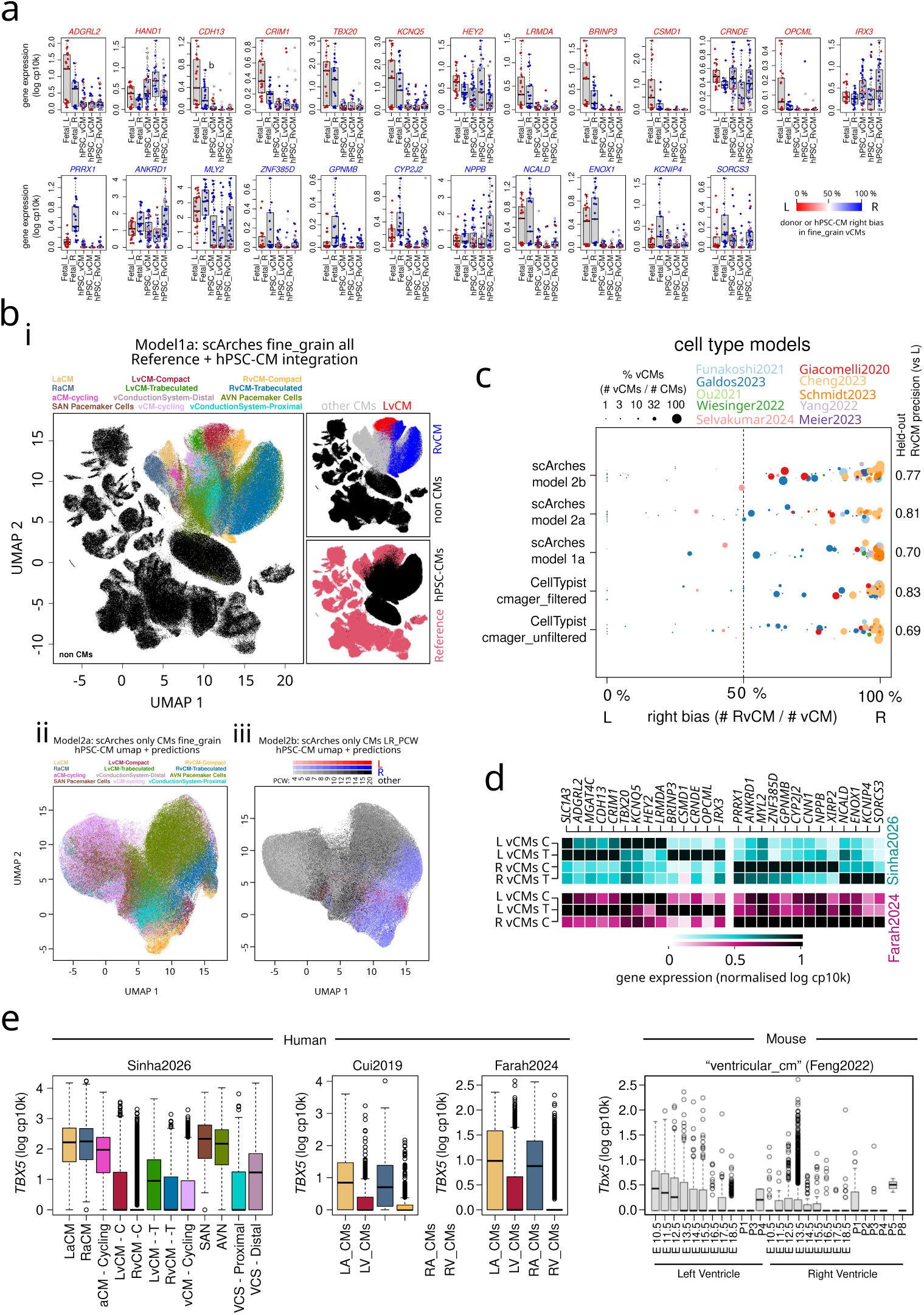
Extended results on left right bias and alternative models. **A:** Gene expression of left (L) and right (R) ventricular cardiomyocyte (vCM) markers in fetal LvCM and fetal RvCM (*n = 21*) across individual donors, and from predicted vCM mid grain, or the subsets LvCM and RvCM fine grain cells from stem cell derived cardiomycyte (hPSC-CM) models (83 conditions, 10 publications). Predictions were filtered (see methods) to remove low confidence classification results (coarse grain CM probability < 0.2). **B:** Three scArches + scANVI models were trained on our dataset shown with their uniform manifold approximation and projection (UMAP) embeddings as i, all data with fine grain annotations in a 63 - class label transfer approach; ii, all cardiomyocytes (CMs) only with a 12-class label transfer model; and iii, CMs as Left or Right explicitly within each donor allowing for across-time confusion between LvCMs or RvCMs. **C:** Right-bias plot across three scArches models, and both filtered and unfiltered `CMageR` CellTypist models trained here in the presented study. **D:** Consistent transcriptional identity of top markers between LvCMs and RvCMs across datasets shown with integrated and re-annotated LvCMs and RvCMs from Farah2024 and this article’s dataset. **E:** Expression of *TBX5* in CM subtypes across three human datasets, and a mouse dataset showing high expression in aCMs, lower in LvCMs, and low but not absent expression in RvCMs.

## 9 Legends for Supplementary Tables

**Supplementary Table 1:** Donor heart metadata for samples used in this study and in Cranley *et al.*[9].

**Supplementary Table 2:** Number of cells within each fine grain annotation across donor, region, and post-conception week.

**Supplementary Table 3:** CMageR CellTypist statistics for coarse, mid, and fine grain models on validation barcodes, and held-out test data.

**Supplementary Table 4:** CMageR CellTypist and alternative scArches model statistics for left vs right ventricular cardiomyocyte (vCM) predictions.

**Supplementary Table 5:** CMageR barcodes and metadata for samples and variable distribution tables used in training of CellTypist and cardiac clock models.

**Supplementary Table 6:** CMageR cardiac clock statistics of RMSE and *R*^2^ for hearts below 15 PCW.

**Supplementary Table 7:** CMageR cardiac clock cardiomyocyte top feature metadata, and individual gene smoother and derivative matrices for the coarse grain cardiomyocyte, and mid grain vCM, aCM, and CCS models ordered by permuted error and feature importance.

**Supplementary Table 8:** Accessed hPSC-CM model datasets and publications information sheet and metadata.

**Supplementary Table 9:** Antibody information sheet.

**Supplementary Table 10:** Highly-variable genes information for scVI.

**Supplementary Table 11:** Differentially expressed cluster markers list for coarse grain cell types.

**Supplementary Table 12:** Differentially expressed cluster markers list for mid grain cell types.

**Supplementary Table 13:** Differentially expressed cluster markers list for fine grain cell types.

